# Potential antiviral options against SARS-CoV-2 infection

**DOI:** 10.1101/2020.05.12.091165

**Authors:** Aleksandr Ianevski, Rouan Yao, Mona Høysæter Fenstad, Svetlana Biza, Eva Zusinaite, Tuuli Reisberg, Hilde Lysvand, Kirsti Løseth, Veslemøy Malm Landsem, Janne Fossum Malmring, Valentyn Oksenych, Sten Even Erlandsen, Per Arne Aas, Lars Hagen, Caroline H. Pettersen, Tanel Tenson, Jan Egil Afset, Svein Arne Nordbø, Magnar Bjørås, Denis E. Kainov

## Abstract

As of June 2020, the number of people infected with severe acute respiratory coronavirus 2 (SARS-CoV-2) continues to skyrocket, with more than 6,5 million cases worldwide. Both the World Health Organization (WHO) and United Nations (UN) has highlighted the need for better control of SARS-CoV-2 infections. However, developing novel virus-specific vaccines, monoclonal antibodies and antiviral drugs against SARS-CoV-2 can be time-consuming and costly. Convalescent sera and safe-in-man broad-spectrum antivirals (BSAAs) are readily available treatment options. Here we developed a neutralization assay using SARS-CoV-2 strain and Vero-E6 cells. We identified most potent sera from recovered patients for treatment of SARS-CoV-2-infected patients. We also screened 136 safe-in-man broad-spectrum antivirals against SARS-CoV-2 infection in Vero-E6 cells and identified nelfinavir, salinomycin, amodiaquine, obatoclax, emetine and homoharringtonine. We found that combinations of virus-directed nelfinavir along with host-directed amodiaquine exhibited the highest synergy. Finally, we developed a website to disseminate the knowledge on available and emerging treatments of COVID-19.

## 1. Introduction

Every year, emerging and re-emerging viruses such as SARS-CoV-2, SARS-CoV, Middle East respiratory syndrome coronavirus (MERS-CoV), Zika virus (ZIKV), Ebola virus (EBOV), influenza A virus (FLUAV) and Rift Valley fever virus (RVFV) surface from natural reservoirs to infect people [1, 2]. As of June 2020, the number of people infected with SARS-CoV-2 continues to rise, with more than 6,5 million cases worldwide.

Coronaviruses (CoV) are a broad family of viruses, which include species such as SARS-CoV-2, SARS-CoV, MERS-CoV, HCoV-229E, HCoV-OC43, HCoV-NL63 and HCoV-HKU1. CoV virions are composed of single-stranded positive-sense RNA, a lipid envelope, and several proteins, including the spike (S), envelope (E), membrane (M), and nucleocapsid (N). HCoV-229E, HCoV-OC43, HCoV-NL63 and HCoV-HKU1 are usually associated with mild, self-limiting upper respiratory tract infections like the common cold. By contrast, SARS-CoV-2, MERS-CoV or SARS-CoV infections can lead to serious disease and death. Considering the current global pandemic, both the WHO and UN has highlighted the need for better control of SARS-CoV-2 infections; however, developing novel virus-specific vaccines, monoclonal antibodies and antiviral drugs against SARS-CoV-2 can be time-consuming and costly [3–10]. Convalescent plasma and safe-in-man broad-spectrum antivirals (BSAAs) are readily available treatment options that can circumvent these difficulties.

Human convalescent plasma is collected from recently recovered individuals and is used to transfer passive antibody immunity to those who have recently been infected or have yet to be exposed to the virus. However, the reliability of diagnostic assays, as well as the quantity and neutralizing capacity of antibodies in the plasma should be considered. Both the WHO and European Centre for Disease Prevention and Control (ECDC) have recommended the evaluation of convalescent plasma for prevention and treatment of COVID-19 disease in controlled trials (WHO/HEO/R&D Blueprint (nCoV)/2020.1) [11]. Indeed, several trials have shown that convalescent plasma reduced viral load and was safe for the treatment of patients with COVID-19 [12, 13]. Interestingly, the studies also revealed variability in specific antibody production that was related to the degree of symptomatic disease. Typically, IgM and IgG antibodies develop between 6–15 days after disease onset, with the seroprevalence increasing to near 100% within 3 weeks. Some recovered patients did not have high titers of neutralizing antibodies, possibly suggesting that antibody levels decline with time and/or that only a subset of patients produces high-titer responses. It is also possible that non-neutralizing antibodies and cellular responses not connected to antibody production are contributing to protection and recovery, as described for other viral diseases [14–16].

BSAAs are small molecules that inhibit human viruses belonging to 2 or more viral families, and that have passed the first phase of clinical trials. We have recently reviewed and summarized the information on dozens of safe-in-man BSAAs in the freely accessible database (https://drugvirus.info/)[17–19]. Forty-six of these agents inhibited SARS-CoV, MERS-CoV, HCoV-229E, HCoV-OC43, HCoV-NL63 and/or HCoV-HKU1. Additionally, clinical investigations have started recently into the effectiveness of BSAAs such as lopinavir, ritonavir, remdesivir, hydroxychloroquine and arbidol against COVID-19 (NCT04252664; NCT0425487; NCT04255017; NCT04261517, NCT04260594). Such BSAAs represent a promising source of drug candidates for SARS-CoV-2 prophylaxis and treatment.

Here, we report isolation of seven different SARS-CoV-2 strains from COVID-19 patients in Norway. We showed that UVC and high temperatures destroyed SARS-CoV-2, establishing a rationale and methodology for safe work in the laboratory. We developed a neutralization assay using SARS-CoV-2 virus and monkey Vero-E6 cells and identified most potent sera from recovered patients for treatment of SARS-CoV-2-infected patients. We also screened 136 safe-in-man broad-spectrum antivirals and identified anti-SARS-CoV-2 activity of salinomycin, nelfinavir, amodiaquine, obatoclax, emetine and homoharringtonine *in vitro*. We further showed that combinations of virus-directed drug nelfinavir and host-directed drugs, such as salinomycin, amodiaquine, obatoclax, emetine and homoharringtonine act synergistically. Finally, we developed a website summarizing scientific and clinical information regarding available and emerging anti-SARS-CoV-2 options.

## 2. Materials and Methods

### 2.1. Patient samples

We selected subjects among healthy individuals, in- and outpatients, as well as patients recovered from SARS-CoV-2 or endemic coronavirus infections. Hospitalization was determined by whether a patient was able to manage symptoms effectively at home, according to local guidelines. ICU admittance was evaluated consistently with the WHO interim guidance on “Clinical management of severe acute respiratory infection when COVID-19 is suspected” (WHO/2019-nCoV/clinical/2020.4). The patients gave their informed consent through the koronastudien.no website. For ICU patients, consent for sample collection was received after treatment or from relatives. For children, consent for sample collection was received from their parents. Donors were recruited through information at the blood collection centres websites and through national media. Nasopharyngeal swabs (NPS) and blood samples were collected. All patients were treated in accordance with good clinical practice, following study protocols. The study was approved by national ethical committee (clinical trial: NCT04320732; REK: 124170).

### 2.1. Cell cultures

Human telomerase reverse transcriptase-immortalized retinal pigment epithelial RPE, lung adenocarcinoma A427, non-small-cell lung cancer Calu-3 and epithelial colorectal adenocarcinoma Caco-2 cells were grown in DMEM-F12 medium supplemented with 100 μg/mL streptomycin and 100 U/mL penicillin (Pen/Strep), 2 mM L-glutamine, 10% FBS, and 0.25% sodium bicarbonate (Sigma-Aldrich, St. Louis, USA). Human neural progenitor cells derived from iPS cells were generated and maintained as described previously [74]. Human large-cell lung carcinoma NCI-H460, colon cancer SW620, colorectal carcinoma SW480 and HT29 cells were grown in RPMI medium supplied with 10% FBS and Pen-Strep. Human adenocarcinoma alveolar basal epithelial A549, human embryonic kidney HEK-293 cells and kidney epithelial cells extracted from an African green monkey (Vero-E6) were grown in DMEM medium supplemented with 10% FBS and Pen-Strep. The cell lines were maintained at 37°C with 5% CO_2_.

### 2.2. Virus isolation

The SARS-CoV-2 strains were isolated and propagated in a Biological Safety Level 3 (BSL-3) facility. 200 μl of nasopharyngeal swabs (NPS) in universal transport medium were diluted 5 times in culture medium (DMEM) supplemented with 0.2% bovine serum albumin (BSA), 0.6 μg/mL penicillin, 60 μg/mL streptomycin, and 2 mM L-glutamine and inoculated into Vero E6 cells. After 4 days of incubation, the media were collected, and the viruses were passaged once again in Vero E6 cells. After 3 days a clear cytopathic effect (CPE) was detected, and virus culture was harvested. Virus concentration was determined by RT-qPCR and plaque assays.

### 2.3. Virus detection and quantification

Viral RNA was extracted using the NTNU_MAG_V2 protocol, a modified version of the BOMB-protocol [75]. 2,5 or 5 μL of the eluate were analysed by RT-qPCR using CFX96 Real-Time Thermal Cycler (Bio-Rad, Hercules, California, USA) as described elsewhere [76] with some modifications. One 20 μl reaction contained 10 μl of QScript XLT One-Step RT-qPCR ToughMix (2x) (Quanta BioSciences, USA), 1 μl each of primer and probe with final respective concentrations of 0,6 μM and 0,25 μM, and 2 μl of molecular grade water. Thermal cycling was performed at 50 °C for 10 min for reverse transcription, followed by 95 °C for 1 min and then 40 cycles of 95 °C for 3 s, 60 °C for 30 s.

For testing the production of infectious virions, we titered the viruses as described in our previous studies [77–79]. In summary, media from the viral culture were serially diluted from 10^−2^ to 10^−7^ in serum-free DMEM media containing 0.2% bovine serum albumin. The dilutions were applied to a monolayer of Vero-E6 cells in 6 or 24-well plates. After one hour, cells were overlaid with virus growth medium containing 1% carboxymethyl cellulose and incubated for 96 h. The cells were fixed and stained with crystal violet dye, and the plaques were calculated in each well, expressed as plaque-forming units per mL (pfu/mL).

### 2.4. Viral genome sequencing

Viral RNA was extracted using the NTNU_MAG_V2 protocol, a modified version of the BOMB-protocol [2]. Libraries were prepared using NuGEN Trio RNA-seq kit. Sequencing was performed on NextSeq500 instrument (NS500528; set up: PE 2 × 75 bp + single index (8 bp)) using NextSeq MID150 sequencing kit, and NextSeq MID flowcell (NCS version: 2.2.0.4). Reads were aligned using the Bowtie 2 software package version 2.3.4.1 to the reference viral genome Wuhan-Hu-1/2019. Sequence alignments were converted to Binary alignments and sorted using SAMtools version 1.5. The consensus FASTQ sequences were obtained with bcftools and vcfutils.pl (from SAMtools) and converted to FASTA using seqtk (https://github.com/lh3/seqtk). Viral genomes in FASTA formats were submitted to www.gisaid.org. The accession numbers of these genomes are: EPI_ISL_450352 (hCoV-19/Norway/Trondheim-E10/2020), EPI_ISL_450351 (hCoV-19/Norway/Trondheim-E9/2020), EPI_ISL_450350 (hCoV-19/Norway/Trondheim-S15/2020), EPI_ISL_450349 (hCoV-19/Norway/Trondheim-S12/2020), EPI_ISL_450348 (hCoV-19/Norway/Trondheim-S10/2020), EPI_ISL_450347 (hCoV-19/Norway/Trondheim-S5/2020), EPI_ISL_450346 (hCoV-19/Norway/Trondheim-S4/2020). Transmission of the strains was visualized using https://nextstrain.org/ncov/global?f_author=Aleksandr%20Ianevski%20et%20al.

### 2.5. UVC- and thermostability assays

The virus (moi 0.1) was aliquoted in Eppendorf tubes and incubated at −80, −20, 4, 20, 37, and 50°C for 48 h or at 96°C for 10 min. Alternatively, the virus was aliquoted in wells of 96 well plate without a lid. The virus was exposed to UVC (λ = 254 nm, ≥ 125 μW/cm^2^) for 10, 20, 40, 80, 160, 320 and 640 s using a germicidal lamp in a biosafety cabinet. The thermo- and UVC-stability of viral RNA was analysed by RT-qPCR. Subsequently, Vero-E6 cells were infected with the virus for 72 h, and cell viability was measured using a CellTiter-Glo assay (Promega, Madison, USA). Luminescence was read using a plate reader.

### 2.6. Neutralization assay

Approximately 4 × 10^4^ Vero-E6 cells were seeded per well in 96-well plates. The cells were grown for 24 h in DMEM medium supplemented with 10% FBS, and Pen/Strep. Serum samples were diluted 100 folds and protein concentrations were quantified using NanoDrop. Serum samples were prepared in 3-fold dilutions at 7 different concentrations starting from 1 mg/mL in the virus growth medium (VGM) containing 0.2% BSA, PenStrep in DMEM. Virus hCoV-19/Norway/Trondheim-E9/2020 was added to the samples to achieve an moi of 0.1 and incubated for 1h at 37 °C. No plasma was added to the control wells. The Vero-E6 Cells were incubated for 72 h with VGM. After the incubation period, the medium was removed and a CellTiter-Glo assay was performed to measure viability.

### 2.7. Drug test

We have previously published a library of safe-in-man BSAAs [5]. Table S1 lists these and other potential BSAAs, their suppliers and catalogue numbers. To obtain 10 mM stock solutions, compounds were dissolved in dimethyl sulfoxide (DMSO, Sigma-Aldrich, Germany) or milli-Q water. The solutions were stored at −80 °C until use.

Approximately 4 × 10^4^ Vero-E6 cells were seeded per well in 96-well plates. The cells were grown for 24h in DMEM medium supplemented with 10% FBS, and Pen/Strep. The medium was replaced with VGM containing 0.2% BSA, and PenStrep in DMEM. The compounds were added to the cells in 3-fold dilutions at 7 or 8 different concentrations starting from 30 μM. No compounds were added to the control wells. The cells were mock- or hCoV-19/Norway/Trondheim-E9/2020-infected at an moi of 0.1. After 72 h of infection, the medium was removed from the cells and a CellTiter-Glo assay was performed to measure viability.

### 2.8. Drug and serum sensitivity quantification

The half-maximal cytotoxic concentration (CC_50_) for each compound was calculated based on viability/death curves obtained on mock-infected cells after non-linear regression analysis with a variable slope using GraphPad Prism software version 7.0a (GraphPad Software, CA, USA). The half-maximal effective concentrations (EC_50_) were calculated based on the analysis of the viability of infected cells by fitting drug dose–response curves using four-parameter (*4PL*) logistic function *f(x)*:

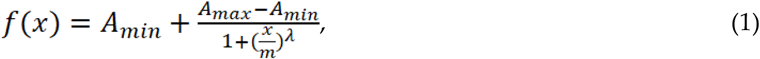

where *f(x)* is a response value at dose *x*, *A*_*min*_ and *A*_*max*_ are the upper and lower asymptotes (minimal and maximal drug effects), *m* is the dose that produces the half-maximal effect (EC_50_ or CC_50_), and *λ* is the steepness (slope) of the curve. A relative effectiveness of the drug was defined as selectivity index (*SI* = CC_50_/EC_50_). The threshold of SI used to differentiate between active and inactive compounds was set to 3.

Area under dose-response curve *AUC* was quantified as:

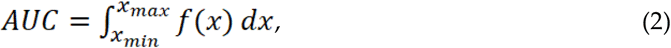

using numerical integration implemented in MESS R package (Bell Laboratories, USA), where *x*_*max*_ and *x*_*min*_ are the maximal and minimal measured doses. Serum sensitivity score (*SSS*) was quantified as a normalized version of standard *AUC* (with the baseline noise subtracted, and normalization of maximal response at the highest concentration often corresponding to off-target toxicity) as

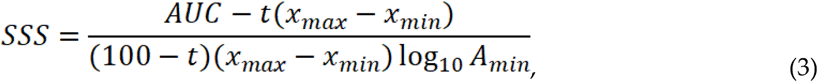

where activity threshold *t* equals 10%.

### 2.9. Drug combination test and synergy calculations

Cells were treated with different concentrations of a combination of two BSAAs. After 72 h, cell viability was measured using a CellTiter-Glo assay. To test whether the drug combinations act synergistically, the observed responses were compared with expected combination responses. The expected responses were calculated based on ZIP reference model using SynergyFinder web-application, version 2 [80].

### 2.10. ELISA assays

We measured the IgG and IgM in human serum using Epitope Diagnostics enzyme linked immunosorbent assays (ELISA) according to manufacturer specifications (Epitope Diagnostics, San Diego, CA). Background-corrected OD values were divided by the cutoff to generate signal-to-cutoff (s/co) ratios. Samples with s/co values greater than 1.0 were considered positive. The Pearson correlation coefficients were calculated by means of stats R package, with the significance determined using Student’s t test.

### 2.11. https://sars-coronavirus-2.info/ website

We reviewed the current landscape of available diagnostic tools, as well as emerging treatment and prophylactic options for the SARS-CoV-2 pandemic and have summarized the information in a database that can be freely accessed at https://sars-coronavirus-2.info. The information in the database was obtained from PubMed, clinicaltrials.gov, DrugBank, DrugCentral, Chinese Clinical Trials Register (ChiCTR), and EU Clinical Trials Register databases [25–27] as well as other public sources. The website was developed with PHP v7 technology using D3.js v5 (https://d3js.org/) for visualization. The COVID-19 statistics is automatically exported from COVID-19 Data Repository by the Center for Systems Science and Engineering (CSSE) at Johns Hopkins University (https://github.com/CSSEGISandData/COVID-19).

## 3. Results

### 3.1. Isolation of SARS-CoV-2 from COVID-19 patients

We isolated 7 SARS-CoV-2 strains from 22 NPS samples of COVID-19 patients using Vero-E6 cells. The RT-qPCR cycle threshold (Ct) values were 18-20 before and 13-15 after propagation of the viruses in Vero-E6 cells (Fig. 1a,b). We sequenced 7 strains and found that the sequences differed from reference hCoV-19/Wuhan/WIV04/2019 strain by a few missense mutations (Fig. 1c). Phylogenetic analysis showed closed relationship between the strains (Fig. 1d). In cross-referencing our sequence data with the pathogen tracking resource, NextStrain.org, we determined that the SARS-CoV-2 strains isolated in Trondheim had originated from China, Denmark, USA and Canada (Fig. 1e).

**Figure 1.**
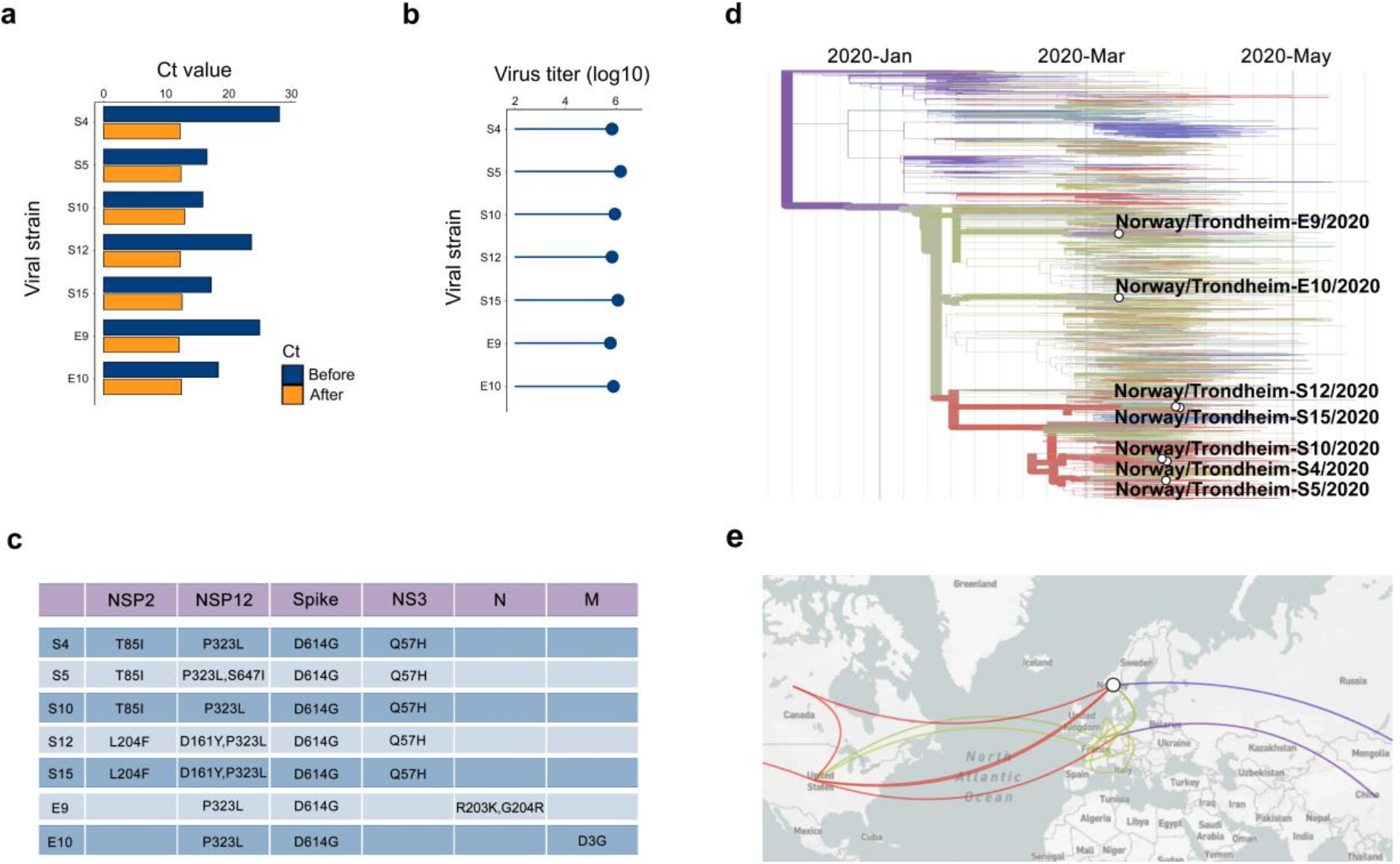
Isolation and characterization of 7 SARS-CoV-2 strains used in this study. (a) RT-qPCR analysis of 7 SARS-CoV-2 strains isolated from nasopharyngeal swabs. (b) Viruses amplified in Vero-E6 cells were quantified by plaque assay. (c) Table showing variations in amino acids between our SARS-CoV-2 strains and the reference hCoV-19/Wuhan/WIV04/2019 strain. (d) Phylogenetic analysis of 7 SARS-CoV-2 strains from Trondheim and other viral strains, which full-genome sequences were submitted to GISAID database. (e) The origin of our 7 strains according to nextstrain.org.

In addition, we tested several cell lines and found that Vero-E6 was the most susceptible for virus-mediated death and virus amplification (Fig. S1a-c). To establish the rationale for safe work, we incubated the virus at different temperatures for 48 h or exposed to UVC radiation for different time periods. The resulting virus preparations were analyzed by RT-qPCR and used to infect Vero-E6 cells. Virus incubation at 37°C for 48 h or UVC exposure for 10 sec destabilized the virus and rescued infected cells from virus-mediated death (Fig. S2).

### 3.2. Sera from patients recovered from COVID-19 contain antibodies which neutralize SARS-CoV-2

We evaluated neutralization capacity of 7 serum samples obtained from patients recovered from COVID-19. We also used sera from patients recovered from endemic coronavirus infections and from healthy blood donors as controls. Five samples from patients recovered from COVID-19 had serum sensitivity score (SSS) > 5 and rescued > 50% cells from virus-mediated death at 1 mg/mL (Fig. 2a,b). Notably, that serum from recovered patient with SSS = 7,2 neutralized all 7 SARS-CoV-2 strains (Fig. S3).

**Figure 2.**
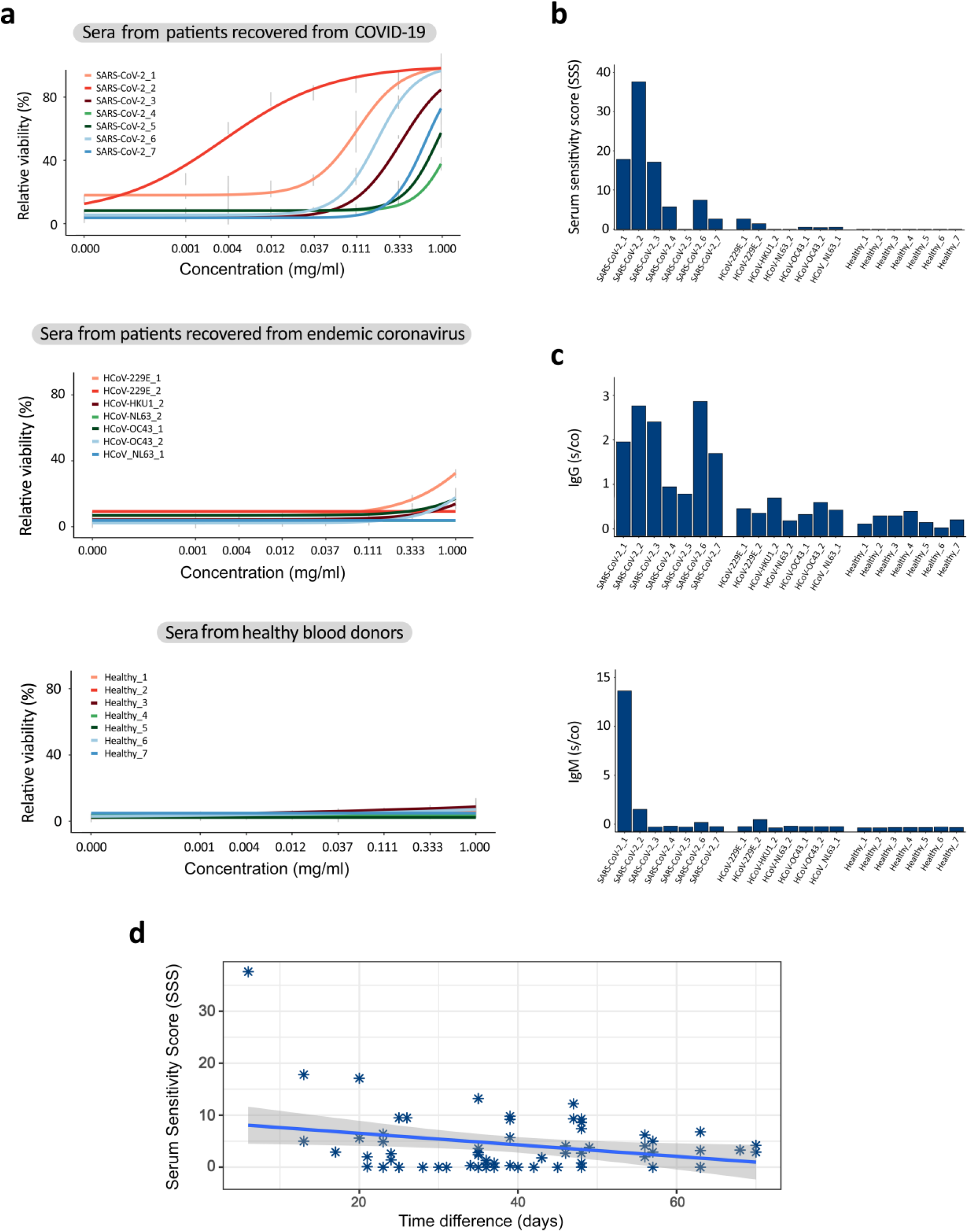
Sera from patients recovered from COVID-19 neutralized SARS-CoV-2 virus and prevented virus-mediated death of Vero-E6 cells. (a) HCoV-19/Norway/Trondheim-E9/2020 strain (moi 0,1) was incubated with indicated concentrations of sera obtained from 7 patients recovered from COVID-19, 7 patients recovered from endemic coronavirus infections and 7 healthy blood donors. The mixtures were added to Vero-E6 cells. Cell viability was measured after 72 h. Mean ± SD, n = 3. (b) Serum sensitivity scores (SSS) were calculated based on curves in (a). (b,c) The IgG and IgM levels were analyzed in the sera of these patients using commercial Elisa kits. (d) Correlation analysis of serum sensitivity scores and time intervals between SARS-CoV-2 diagnosis and sera collection.

Our neutralization test of 32 samples (Table S2) showed a moderate positive correlation with IgG (r=0.59, p < 0.001) and IgM (r=0.43, p = 0.01) s/co values obtained using commercial ELISA kits that recognize SARS-CoV-2 N protein (Fig. 2c,d). However, correlation between the IgG and IgM ELISA results was only r=0.28, p=0.11. Furthermore, we found moderate negative correlation between SSSs and time intervals between SARS-CoV-2 diagnosis and serum collection for 66 samples (−0.5, p < 0.025; Fig. 2d). Altogether, these results suggest that patients diagnosed with COVID-19 produce different immune responses to SARS-CoV-2 infection and that neutralization capacity of convalescent sera declines with time.

### 3.3. Repurposing safe-in-man BSAAs

Through literature review, we made a database to summarize safe-in-man BSAAs (https://drugvirus.info/). Recently, we have expanded on the spectrum of activities for some of these agents [18, 19, 28–30]. Some of these agents could be repositioned for treatment of SARS-CoV-2 infection.

We tested 136 agents against SARS-CoV-2 in VERO-E6 cells. Remdesivir was included as a positive control [31], and nicotine as a negative control. Seven different concentrations of the compounds were added to virus-infected cells. Cell viability was measured after 72 h to determine compound efficiency. After the initial screening, we identified apilimod, emetine, amodiaquine, obatoclax, homoharringtonine, salinomycin, arbidol, posaconazole and nelfinavir as compounds that rescued virus-infected cells from death (AUC from 285 to 585; Table S1). The compounds we identified possessed structure-activity relationship (Fig. 3a). AUC for remdesivir was 290. Interestingly, 10 μM nicotine rescued cells from virus-mediated death but altered cell morphology (AUC = 239; Fig. S4).

**Figure 3.**
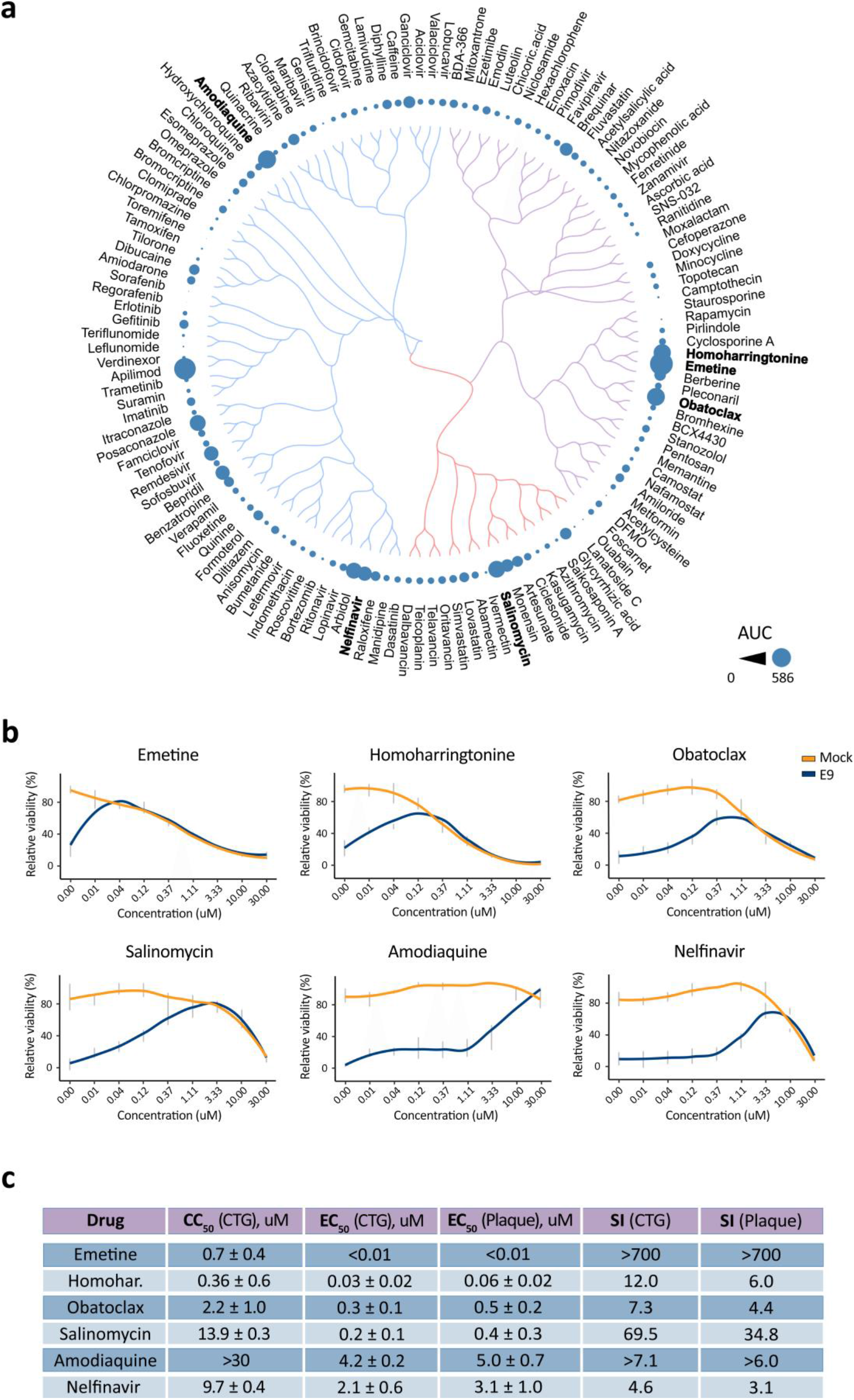
Anti-SARS-CoV-2 activity of safe-in man broad-spectrum antivirals in Vero-E6 cells. (a) Structure-antiviral activity relation of 136 BSAAs. The compounds were clustered based on their structural similarity calculated by ECPF4 fingerprints and visualized using D3 JavaScript library. The anti-SARS-CoV-2 activity of the compounds was quantified using AUC and shown as bubbles. Bubble size corresponds to compounds AUCs. (b) Vero-E6 cells were treated with increasing concentrations of a compound and infected with HCoV-19/Norway/Trondheim-E9/2020 strain (moi, 0.1) or mock. After 72 h the viability of the cells was determined using the CTG assay. Mean ± SD; n = 3. (c) Table showing half-maximal cytotoxic concentration (CC50), the half-maximal effective concentration (EC50), and selectivity indexes (SI=CC50/EC50) for selected anti-SARS-CoV-2 compounds calculated from CTG and plaque assays. Mean ± SD; n = 3.

We repeated the experiment with hit compounds monitoring their toxicity and efficacy. We confirmed antiviral activity of emetine, amodiaquine, obatoclax, homoharringtonine, salinomycin, and nelfinavir (Fig 3b,c). Importantly, amodiaquine had superior SI over its analogues, chloroquine, hydroxychloroquine, quinacrine, and mefloquine (Fig. S5). Thus, we identified and validated anti-SARS-CoV-2 activities for six BSAAs in Vero-E6 cells.

### 3.4. BSAA combinations are effective against SARS-CoV-2 infection

To test for potential synergism among the validated hits, we treated cells with varying concentrations of a two-drug combination and monitored cell viability (Fig. 4a). The observed drug combination responses were compared with expected combination responses calculated by means of the zero-interaction potency (ZIP) model [24, 32]. We quantified synergy scores, which represent the average excess response due to drug interactions (i.e. 10% of cell survival beyond expected additivity between single drugs has a synergy score of 10). We found that combinations of nelfinavir with salinomycin, amodiaquine, homoharringtonine, and obatoclax, as well as combination of amodiaquine and salinomycin were synergistic (most synergistic area scores >10; Fig. 4b). Moreover, nelfinavir-amodiaquine treatment was effective against all 7 SARS-CoV-2 strains (Fig. 4c). Thus, we identified synergistic combination of nelfinavir-amodiaquine against SARS-CoV-2 infections.

**Figure 4.**
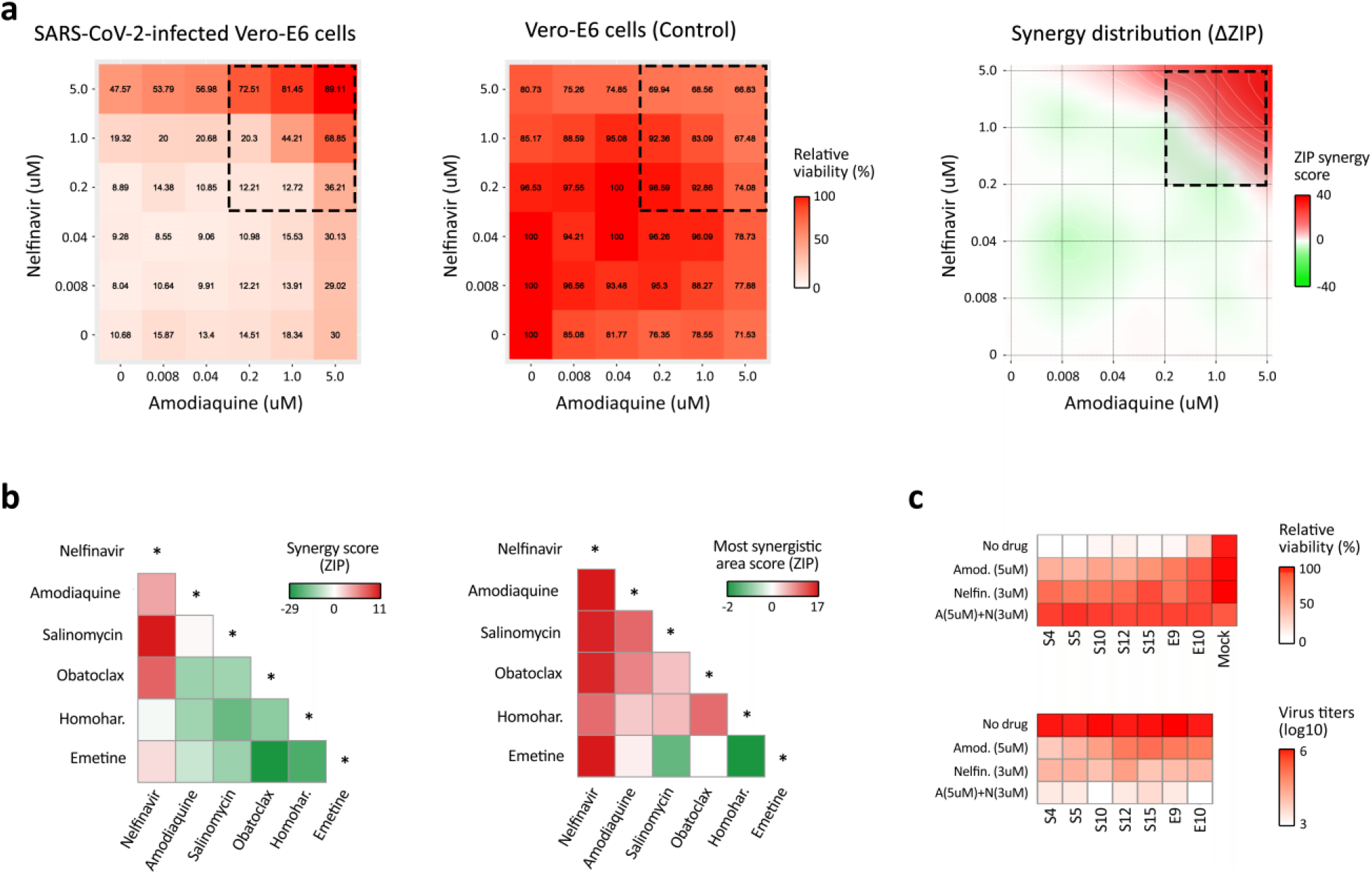
Effect of drug combinations on SARS-CoV-2 infection. (a) The representative interaction landscapes of one of the drug combinations (amodiaquine-nelfinavir) measured using CTG assay and SARS-CoV-2- and mock-infected cells (left and central panels). Synergy distribution is shown for virus-infected cells (right panel). (b) Synergy scores and the most synergistic area scores of 15 drug combinations. (c) The effect of amodiaquine-nelfinavir combination on viral replication (lower panel) and viability of cells infected with 7 different SARS-CoV-2 strains (upper panel).

### 3.5. Sars-coronavirus-2.info website summarizes emerging antiviral options

To rapidly respond to the COVID-19 outbreak, we developed a freely accessible website summarizing novel anti-SARS-CoV-2 developments, including the tracking of therapeutic/antiviral drug development, development of vaccines, and currently approved diagnostic options around the globe.

The ‘Treatment’ section of the website summarizes 542 in-progress and completed clinical trials that test the efficacy of therapeutic agents to treat COVID-19 or complications that arise from COVID-19. These trials include over 192 unique therapeutic agents, in varying combinations and applications. Importantly, we list 71 clinical trials that are already completed or are projected to be completed by the end of June 2020. Of note among these are trials of remdesivir, favipiravir, lopinavir/ritonavir, hydroxychloroquine, dipyridamole, and interferons alpha and beta, which are all phase 3 or 4 clinical trials scheduled to be currently completed.

The ‘Prevention’ section summarizes 23 current vaccine trials taking place around the globe. Although vaccine development lags considerably behind drug development, several repurposed vaccine options have also emerged. This includes trials of the cross-reactivity of the MMR (Measles, Mumps, Rubella) vaccine, as well as several trials of the BCG vaccine among high-risk populations, such as health care workers.

Finally, the ‘Testing’ section of the website provides a summary of 377 currently available laboratory-based and point-of-care diagnostic tests that are approved for clinical diagnosis in at least one country.

The website also includes predictions of experimental and approved drugs effective against SARS-CoV-2, as well as provides a summary of information about the coronavirus pandemic. The website allows interactive exploration of the data with built-in feedback and is available in several languages. The website is updated as soon as novel anti-SARS-CoV-2 options emerge, or the statuses of existing ones are updated.

## 4. Discussion

Here, we reported the isolation of 7 SARS-CoV-2 strains from samples of patients suffering from COVID-19. Full-genome sequencing revealed that the strains were highly similar (98,8%) to one another and to the strains circulating in China, Denmark, USA and Canada. All 7 strains contain D614G in S protein. Strains with this mutation began spreading in Europe in early February and became the dominant in other regions (https://doi.org/10.1101/2020.04.29.069054).

We screened 12 cell lines for their susceptibility for SARS-CoV-2 infection and virus replication. Vero-E6 cells appeared to be the most susceptible cell line for virus-mediated cell death and virus propagation. This cell line has been widely used on toxicology, virology and pharmacology research, as well as, on the production of vaccines and diagnostic reagents. The cell line is interferon-deficient; unlike normal mammalian cells, it does not secrete interferon alpha or beta when infected by viruses [81]. Moreover, this cell line was used routinely in anti-SARS-CoV-2 research [82–85].

We also showed that >10 sec of UVC radiation or 48 h incubation at 37 °C neutralized SARS-CoV-2, establishing a rationale and methodology for safe work in the laboratory. These results are consistent with previous studies showing that physical factors destabilize SARS-CoV-2 and other viruses [33–36].

Neutralization tests are crucial tools for the assessment of previous SARS-CoV-2 exposure and potential immunity [12, 13]. We have developed a test to assess the neutralization capacity of serum samples from patients recovered from SARS-CoV-2 infections, patients with endemic coronavirus infections and healthy blood donors. Our results suggest that COVID-19 patients respond differently to SARS-CoV-2 infection. Moreover, neutralization capacity of convalescence sera declined with time. Thus, the neutralization test allowed us to identify the most potent sera from patients recovered from COVID-19 for treatment of SARS-CoV-2-infected patients.

Moreover, results from our neutralization test positively correlated with those from commercial ELISA assays. However, correlation between the IgG and IgM ELISAs results was only moderate. The difference could be associated with time of sample collection, production of immunoglobulins, or sensitivity that can be attributed to the technique and the antigen in use (i.e., IgM is the first immunoglobulin to be produced in response to an antigen and can be detected during early onset of disease, whereas IgG is maintained in the body after initial exposure for long term response and can be detected after the infection has passed).

Drug repurposing, also called drug repositioning, is a strategy for generating additional value from an existing drug by targeting disease other than that for which it was originally intended [37, 38]. This has significant advantages over new drug discovery since chemical synthesis steps, manufacturing processes, reliable safety, and pharmacokinetic properties in pre-clinical (animal model) and early clinical developmental phases (phase 0, I and IIa) are already. Therefore, drug repositioning for COVID-19 provides unique translational opportunities, including a substantially higher probability of success to market as compared with developing new virus-specific drugs and vaccines, as well as significantly reduced cost and timeline to clinical availability [30, 39, 40].

We tested 136 safe-in-man BSAAs against SARS-CoV-2 in cell culture. We identified salinomycin, obatoclax, amodiaquine, nelfinavir, emetine, and homoharringtonine as having anti-SARS-CoV-2 activity, which we put forward as potential anti-SARS-CoV-2 drug candidates.

Nelfinavir (Viracept) is an orally bioavailable inhibitor of human immunodeficiency virus HIV-1 (750 mg PO q8hr). It targets HIV protease for the treatment of HIV infection [41]. Molecular docking studies predict that nelfinavir binds to the SARS-CoV-2 protease [42]. Nelfinavir could also inhibit cell fusion caused by the SARS-CoV-2 S glycoprotein (https://doi.org/10.1101/2020.04.06.026476) [43]. It also inhibits Chikungunya virus (CHIKV), Dengue virus (DENV), hepatitis C virus (HCV), herpes simplex virus 1 (HSV-1), and SARS-CoV infections (https://doi.org/10.1039/C5RA14469H) [44–46]. These studies warrant further investigations of the potential of nelfinavir alone or in combination with other drugs to inhibit SARS-CoV-2 infections.

Amodiaquine is a medication used to treat malaria. The recommended dose for a course of amodiaquine is 30 mg amodiaquine base/kg body weight over 3 days, i.e., 10 mg/kg/day [86]. It also shows broad-spectrum activity against ZIKV, DENV, HCV, MERS-CoV, SARS-CoV, SARS-CoV-2, Ross River virus (RRV), sindbis virus (SINV), West Nile virus (WNV), yellow fever virus (YFV), EBOV, Lassa virus (LASV), rabies virus (RABV), Vesicular Stomatitis Virus (VSV) and HSV-1 viruses (https://doi.org/10.1101/2020.03.25.008482) [51–56]. Importantly, amodiaquine showed more potent antiviral activity than its analogues chloroquine and hydroxychloroquine. A recent study showed that hydroxychloroquine and chloroquine have ‘no benefit’ for coronavirus patients and could even increase the risk of heart arrhythmias and mortality (https://doi.org/10.1016/S0140-6736(20)31180-6).

Obatoclax was originally developed as an anticancer agent. Several phase II clinical trials have investigated the use of obatoclax in the treatment of leukemia, lymphoma, myelofibrosis, and mastocytosis. Continuous 24-hour infusion of obatoclax 25-60 mg/day for 3 days in 2-week cycles or 3-hour infusion in 3 d cycle have previously been evaluated in cancer patients [87]. In addition, obatoclax showed antiviral activity against FLUAV, ZIKV, WNV, YFV, SINV, Junín Virus (JUNV), LASV, herpes simplex virus 2 (HSV-2), echovirus 1 (EV1), human metapneumovirus (HMPV), RVFV and lymphocytic choriomeningitis virus (LCMV) in vitro [18, 57–60]. It was shown that obatoclax inhibited viral endocytic uptake by targeting cellular Mcl-1 protein [60].

Emetine is an antiprotozoal drug. It is administered by intramuscular or deep subcutaneous injection in a dose of 1 mg/kg/day (maximum, 60 mg/day) for 10 days [88]. Emetine is also used to induce vomiting. In addition, it possesses antiviral effects against ZIKV, EBOV, RABV, cytomegalovirus (CMV), HCoV-OC43, HSV-2, EV1, HMPV, RVFV, FLUAV, HIV-1 and SARS-CoV-2 [18, 61–66] [89] (https://doi.org/10.1101/2020.03.25.008482). Emetine was proposed to inhibit viral polymerases, though it could have some other targets [67].

Homoharringtonine is an anticancer drug which is indicated for treatment of chronic myeloid leukaemia (2 mg/m^2^ IV daily × 7). It also possesses antiviral activities against hepatitis B virus (HBV), MERS-CoV, HSV-1, EV1, VZV and SARS-CoV-2 in vitro [18, 66, 68–71]. Homoharringtonine binds to the 80S ribosome and inhibits viral protein synthesis by interfering with chain elongation [69].

Salinomycin is an orally bioavailable antibiotic, which is used against Gram-positive bacteria in animal husbandry (0.2 mg/kg BW, PO). It also inhibits FLUAV, Respiratory syncytial virus (RSV) and CMV infections [47, 48]. Salinomycin was proposed to disrupt endosomal acidification and to block entry of the viruses into cells [49][50].

Our results have uncovered several existing BSAAs that could be re-positioned to SARS-CoV-2 infections. Because PK/PD and toxicology studies have already been performed on these compounds, *in vivo* efficacy studies could be initiated immediately, saving time and resources.

Combination therapies became a standard for the treatment of HIV and HCV infections. The reasons for using combinations rather than single antiviral are better efficacy, decreased toxicity, and prevention of resistance emergence. Here we found that combinations of nelfinavir with salinomycin, amodiaquine, obatoclax, emetine or homoharringtonine were synergistic against SARS-CoV-2 in Vero-E6 cells. Thus, the synergy was achieved when virus-directed drug was combined with host-directed ones. This observation agrees with other studies on such combinations [18, 58, 72, 73] (https://doi.org/10.1101/2020.04.14.039925).

According to the available pharmacological data for these drugs, the most potent combination could be a combination of orally available nelfinavir and amodiaquine. This was also the combination that exhibited the highest synergy of all the drug combinations we tested, with the synergy score at the most synergistic area being 16.48. (i.e. 16.48% of cell survival beyond expected additivity between single drugs). There are no guidelines of what is considered a good synergy, but it is very common to consider synergy > 10 as true (significant) synergy. Thus, amodiaquine and nelfinavir combination could result in better efficacy and decreased toxicity for the treatment of SARS-CoV-2 and perhaps other viral infections.

Our future goal is to complete preclinical studies and translate our findings into trials in patients. The most effective and tolerable BSAAs or their combinations will have a global impact, improving protection of the general population from emerging and re-emerging viral infections or co-infections, and allowing the rapid management of drug-resistant strains. Our bigger ambition is to assemble a toolbox of BSAAs for the treatment of emerging and re-emerging viral infections. This toolbox can be offered to the WHO as a means for the fast identification of safe and effective antiviral options.

We have summarized the information about the status of currently available and emerging anti-SARS-CoV-2 options in the freely accessible website (https://sars-coronavirus-2.info). The website is updated regularly and incorporates novel anti-SARS-CoV-2 options as they emerge, or the changes the statuses of existing ones as updates occur.

## 5. Conclusions

In conclusion, sera from recovered patients, BSAAs, and combinations of BSAAS as well as other available and emerging treatments could have a pivotal role in the battle against COVID-19 and other emerging and re-emerging viral diseases. Further development of these options could save time and resources that are required for the development of alternatives - virus-specific drugs and vaccines. This could have a global impact by decreasing morbidity and mortality, maximizing the number of healthy life years, improving the quality of life of infected patients, and decreasing the costs of patient care curtailing to the impact of the current SARS-CoV-2 pandemic as well as future viral outbreaks.

## 6. Patents

N.A.

## Supplementary Materials

Table S1: Compounds, their suppliers, catalogue numbers and AUCs; Table S2: Results of neutralization and ELISA assays; Figure S1: Propagation of HCoV-19/Norway/Trondheim-E9/2020 in cell cultures; Figure S2: Effect of temperature and UV radiation on infectivity of HCoV-19/Norway/Trondheim-E9/2020 strain; Figure S3: The effect of serum from patient recovered from SARS-CoV-2 infection on viability of Vero-E6 cells infected with 7 SARS-CoV-2 strains; Figure S4: Comparison of anti-SARS-CoV-2 activities of amodiaquine and its analogues.

## Author Contributions

All authors contributed to methodology, software, validation, formal analysis, investigation, resources, data curation, writing, review or editing of the manuscript. D.K. conceptualized, supervised and administrated the study, and acquired funding. All authors have read and agreed to the published version of the manuscript.

## Acknowledgments

This study was devoted to Li Wenliang, a Chinese doctor who tried to warn about Coronavirus, as well as to many other doctors and COVID-19 patients. We thank Koit Aasumets, Sergio Miguel Castañeda Zegarra, Qindong Zhang, Simona Komarova, Nikki Upfold, Miriam Khider, Hege Hval, and Kasia Kolasa for translation of the sars-coronavirus-2.info website to different languages. We also thank Maxim Bespalov, Sergei Shiryaev, Pavel Uvarov, Evgeny Kulesskiy and many other people for sharing their ideas on drug candidates. In addition, we thank Wei Wang, Marit Bugge and Nadra J. Nilsen for cell lines.

## References

1. WHO. WHO publishes list of top emerging diseases likely to cause major epidemics. wwwwhoint/medicines/ebola-treatment/WHO-list-of-top-emerging-diseases/en/. 2015.

2. Howard CR, Fletcher NF. Emerging virus diseases: can we ever expect the unexpected? Emerg Microbes Infect. 2012;1(12):e46.

3. Bekerman E, Einav S. Infectious disease. Combating emerging viral threats. Science. 2015;348(6232):282–3.

4. Yuan S, Chu H, Chan JF, Ye ZW, Wen L, Yan B, et al. SREBP-dependent lipidomic reprogramming as a broad-spectrum antiviral target. Nat Commun. 2019;10(1):120.

5. Check Hayden E. Experimental drugs poised for use in Ebola outbreak. Nature. 2018;557(7706):475–6.

6. Jaishankar D, Yakoub AM, Yadavalli T, Agelidis A, Thakkar N, Hadigal S, et al. An off-target effect of BX795 blocks herpes simplex virus type 1 infection of the eye. Sci Transl Med. 2018;10(428).

7. Schor S, Einav S. Repurposing of Kinase Inhibitors as Broad-Spectrum Antiviral Drugs. DNA Cell Biol. 2018;37(2):63–9.

8. Debing Y, Neyts J, Delang L. The future of antivirals: broad-spectrum inhibitors. Curr Opin Infect Dis. 2015;28(6):596–602.

9. Yu F, Lu L, Du L, Zhu X, Debnath AK, Jiang S. Approaches for identification of HIV-1 entry inhibitors targeting gp41 pocket. Viruses. 2013;5(1):127–49.

10. De Clercq E, Li G. Approved Antiviral Drugs over the Past 50 Years. Clin Microbiol Rev. 2016;29(3):695–747.

11. Casadevall A, Pirofski LA. The convalescent sera option for containing COVID-19. J Clin Invest. 2020;130(4):1545–8.

12. Duan K, Liu B, Li C, Zhang H, Yu T, Qu J, et al. Effectiveness of convalescent plasma therapy in severe COVID-19 patients. Proc Natl Acad Sci U S A. 2020;117(17):9490–6.

13. Shen C, Wang Z, Zhao F, Yang Y, Li J, Yuan J, et al. Treatment of 5 Critically Ill Patients With COVID-19 With Convalescent Plasma. JAMA. 2020.

14. Jenks JA, Goodwin ML, Permar SR. The Roles of Host and Viral Antibody Fc Receptors in Herpes Simplex Virus (HSV) and Human Cytomegalovirus (HCMV) Infections and Immunity. Front Immunol. 2019;10:2110.

15. van Erp EA, Luytjes W, Ferwerda G, van Kasteren PB. Fc-Mediated Antibody Effector Functions During Respiratory Syncytial Virus Infection and Disease. Front Immunol. 2019;10:548.

16. Gunn BM, Yu WH, Karim MM, Brannan JM, Herbert AS, Wec AZ, et al. A Role for Fc Function in Therapeutic Monoclonal Antibody-Mediated Protection against Ebola Virus. Cell Host Microbe. 2018;24(2):221–33 e5.

17. Andersen PI, Ianevski A, Lysvand H, Vitkauskiene A, Oksenych V, Bjoras M, et al. Discovery and development of safe-in-man broad-spectrum antiviral agents. Int J Infect Dis. 2020;93:268–76.

18. Andersen PI, Krpina K, Ianevski A, Shtaida N, Jo E, Yang J, et al. Novel Antiviral Activities of Obatoclax, Emetine, Niclosamide, Brequinar, and Homoharringtonine. Viruses. 2019;11(10).

19. Ianevski A, Zusinaite E, Kuivanen S, Strand M, Lysvand H, Teppor M, et al. Novel activities of safe-in-human broad-spectrum antiviral agents. Antiviral Res. 2018;154:174–82.

20. Oberacker P, Stepper P, Bond DM, Hohn S, Focken J, Meyer V, et al. Bio-On-Magnetic-Beads (BOMB): Open platform for high-throughput nucleic acid extraction and manipulation. PLoS Biol. 2019;17(1):e3000107.

21. Corman VM, Landt O, Kaiser M, Molenkamp R, Meijer A, Chu DK, et al. Detection of 2019 novel coronavirus (2019-nCoV) by real-time RT-PCR. Euro Surveill. 2020;25(3).

22. Bosl K, Ianevski A, Than TT, Andersen PI, Kuivanen S, Teppor M, et al. Common Nodes of Virus–Host Interaction Revealed Through an Integrated Network Analysis. Front Immunology. 2019;Accepted.

23. Bulanova D, Ianevski A, Bugai A, Akimov Y, Kuivanen S, Paavilainen H, et al. Antiviral Properties of Chemical Inhibitors of Cellular Anti-Apoptotic Bcl-2 Proteins. Viruses. 2017;9(10).

24. Ianevski A, He L, Aittokallio T, Tang J. SynergyFinder: a web application for analyzing drug combination dose-response matrix data. Bioinformatics. 2017;33(15):2413–5.

25. Ursu O, Holmes J, Bologa CG, Yang JJ, Mathias SL, Stathias V, et al. DrugCentral 2018: an update. Nucleic Acids Res. 2019;47(D1):D963–D70.

26. Kim S, Chen J, Cheng T, Gindulyte A, He J, He S, et al. PubChem 2019 update: improved access to chemical data. Nucleic Acids Res. 2019;47(D1):D1102–D9.

27. Wishart DS, Feunang YD, Guo AC, Lo EJ, Marcu A, Grant JR, et al. DrugBank 5.0: a major update to the DrugBank database for 2018. Nucleic Acids Res. 2018;46(D1):D1074–D82.

28. Andersen PI, Ianevski A, Lysvand H, Vitkauskiene A, Oksenych V, Bjoras M, et al. Discovery and development of safe-in-man broad-spectrum antiviral agents. Int J Infect Dis. 2020.

29. Bosl K, Ianevski A, Than TT, Andersen PI, Kuivanen S, Teppor M, et al. Common Nodes of Virus-Host Interaction Revealed Through an Integrated Network Analysis. Front Immunol. 2019;10:2186.

30. Ianevski A, Andersen PI, Merits A, Bjoras M, Kainov D. Expanding the activity spectrum of antiviral agents. Drug Discov Today. 2019;24(5):1224–8.

31. Yin W, Mao C, Luan X, Shen DD, Shen Q, Su H, et al. Structural basis for inhibition of the RNA-dependent RNA polymerase from SARS-CoV-2 by remdesivir. Science. 2020.

32. Ianevski A, Giri AK, Aittokallio T. SynergyFinder 2.0: visual analytics of multi-drug combination synergies. Nucleic Acids Res. 2020.

33. Demongeot J, Flet-Berliac Y, Seligmann H. Temperature Decreases Spread Parameters of the New COVID-19 Case Dynamics. Biology (Basel). 2020;9(5).

34. Cadnum JL, Li DF, Redmond SN, John AR, Pearlmutter B, Donskey CJ. Effectiveness of Ultraviolet-C Light and a High-Level Disinfection Cabinet for Decontamination of N95 Respirators. Pathog Immun. 2020;5(1):52–67.

35. Sobral MFF, Duarte GB, da Penha Sobral AIG, Marinho MLM, de Souza Melo A. Association between climate variables and global transmission oF SARS-CoV-2. Sci Total Environ. 2020;729:138997.

36. Ianevski A, Zusinaite E, Shtaida N, Kallio-Kokko H, Valkonen M, Kantele A, et al. Low Temperature and Low UV Indexes Correlated with Peaks of Influenza Virus Activity in Northern Europe during 2010(-)2018. Viruses. 2019;11(3).

37. Nishimura Y, Hara H. Editorial: Drug Repositioning: Current Advances and Future Perspectives. Front Pharmacol. 2018;9:1068.

38. Pushpakom S, Iorio F, Eyers PA, Escott KJ, Hopper S, Wells A, et al. Drug repurposing: progress, challenges and recommendations. Nat Rev Drug Discov. 2019;18(1):41–58.

39. Pizzorno A, Padey B, Terrier O, Rosa-Calatrava M. Drug Repurposing Approaches for the Treatment of Influenza Viral Infection: Reviving Old Drugs to Fight Against a Long-Lived Enemy. 2019;10(531).

40. Zheng W, Sun W, Simeonov A. Drug repurposing screens and synergistic drug-combinations for infectious diseases. Br J Pharmacol. 2018;175(2):181–91.

41. Zhang KE, Wu E, Patick AK, Kerr B, Zorbas M, Lankford A, et al. Circulating metabolites of the human immunodeficiency virus protease inhibitor nelfinavir in humans: structural identification, levels in plasma, and antiviral activities. Antimicrob Agents Chemother. 2001;45(4):1086–93.

42. Mothay D, Ramesh KV. Binding site analysis of potential protease inhibitors of COVID-19 using AutoDock. Virusdisease. 2020:1–6.

43. Musarrat F, Chouljenko V, Dahal A, Nabi R, Chouljenko T, Jois SD, et al. The anti-HIV Drug Nelfinavir Mesylate (Viracept) is a Potent Inhibitor of Cell Fusion Caused by the SARS-CoV-2 Spike (S) Glycoprotein Warranting further Evaluation as an Antiviral against COVID-19 infections. J Med Virol. 2020.

44. Kalu NN, Desai PJ, Shirley CM, Gibson W, Dennis PA, Ambinder RF. Nelfinavir inhibits maturation and export of herpes simplex virus 1. J Virol. 2014;88(10):5455–61.

45. Toma S, Yamashiro T, Arakaki S, Shiroma J, Maeshiro T, Hibiya K, et al. Inhibition of intracellular hepatitis C virus replication by nelfinavir and synergistic effect with interferon-alpha. J Viral Hepat. 2009;16(7):506–12.

46. Yamamoto N, Yang R, Yoshinaka Y, Amari S, Nakano T, Cinatl J, et al. HIV protease inhibitor nelfinavir inhibits replication of SARS-associated coronavirus. Biochem Biophys Res Commun. 2004;318(3):719–25.

47. Norris MJ, Malhi M, Duan W, Ouyang H, Granados A, Cen Y, et al. Targeting Intracellular Ion Homeostasis for the Control of Respiratory Syncytial Virus. Am J Respir Cell Mol Biol. 2018;59(6):733–44.

48. Kapoor A, He R, Venkatadri R, Forman M, Arav-Boger R. Wnt modulating agents inhibit human cytomegalovirus replication. Antimicrob Agents Chemother. 2013;57(6):2761–7.

49. Jang Y, Shin JS, Yoon YS, Go YY, Lee HW, Kwon OS, et al. Salinomycin Inhibits Influenza Virus Infection by Disrupting Endosomal Acidification and Viral Matrix Protein 2 Function. J Virol. 2018;92(24).

50. Jeon S, Ko M, Lee J, Choi I, Byun SY, Park S, et al. Identification of antiviral drug candidates against SARS-CoV-2 from FDA-approved drugs. Antimicrob Agents Chemother. 2020.

51. Mazzon M, Ortega-Prieto AM, Imrie D, Luft C, Hess L, Czieso S, et al. Identification of Broad-Spectrum Antiviral Compounds by Targeting Viral Entry. Viruses. 2019;11(2).

52. Hulseberg CE, Feneant L, Szymanska-de Wijs KM, Kessler NP, Nelson EA, Shoemaker CJ, et al. Arbidol and Other Low-Molecular-Weight Drugs That Inhibit Lassa and Ebola Viruses. J Virol. 2019;93(8).

53. Sakurai Y, Sakakibara N, Toyama M, Baba M, Davey RA. Novel amodiaquine derivatives potently inhibit Ebola virus infection. Antiviral Res. 2018;160:175–82.

54. Zhou T, Tan L, Cederquist GY, Fan Y, Hartley BJ, Mukherjee S, et al. High-Content Screening in hPSC-Neural Progenitors Identifies Drug Candidates that Inhibit Zika Virus Infection in Fetal-like Organoids and Adult Brain. Cell Stem Cell. 2017;21(2):274–83 e5.

55. Dyall J, Coleman CM, Hart BJ, Venkataraman T, Holbrook MR, Kindrachuk J, et al. Repurposing of clinically developed drugs for treatment of Middle East respiratory syndrome coronavirus infection. Antimicrob Agents Chemother. 2014;58(8):4885–93.

56. Boonyasuppayakorn S, Reichert ED, Manzano M, Nagarajan K, Padmanabhan R. Amodiaquine, an antimalarial drug, inhibits dengue virus type 2 replication and infectivity. Antiviral Res. 2014;106:125–34.

57. Kim YJ, Cubitt B, Chen E, Hull MV, Chatterjee AK, Cai Y, et al. The ReFRAME library as a comprehensive drug repurposing library to identify mammarenavirus inhibitors. Antiviral Res. 2019;169:104558.

58. Kuivanen S, Bespalov MM, Nandania J, Ianevski A, Velagapudi V, De Brabander JK, et al. Obatoclax, saliphenylhalamide and gemcitabine inhibit Zika virus infection in vitro and differentially affect cellular signaling, transcription and metabolism. Antiviral Res. 2017;139:117–28.

59. Varghese FS, Rausalu K, Hakanen M, Saul S, Kummerer BM, Susi P, et al. Obatoclax Inhibits Alphavirus Membrane Fusion by Neutralizing the Acidic Environment of Endocytic Compartments. Antimicrob Agents Chemother. 2017;61(3).

60. Denisova OV, Kakkola L, Feng L, Stenman J, Nagaraj A, Lampe J, et al. Obatoclax, saliphenylhalamide, and gemcitabine inhibit influenza a virus infection. J Biol Chem. 2012;287(42):35324–32.

61. Shen L, Niu J, Wang C, Huang B, Wang W, Zhu N, et al. High-Throughput Screening and Identification of Potent Broad-Spectrum Inhibitors of Coronaviruses. J Virol. 2019;93(12).

62. MacGibeny MA, Koyuncu OO, Wirblich C, Schnell MJ, Enquist LW. Retrograde axonal transport of rabies virus is unaffected by interferon treatment but blocked by emetine locally in axons. PLoS Pathog. 2018;14(7):e1007188.

63. Yang S, Xu M, Lee EM, Gorshkov K, Shiryaev SA, He S, et al. Emetine inhibits Zika and Ebola virus infections through two molecular mechanisms: inhibiting viral replication and decreasing viral entry. Cell Discov. 2018;4:31.

64. Mukhopadhyay R, Roy S, Venkatadri R, Su YP, Ye W, Barnaeva E, et al. Efficacy and Mechanism of Action of Low Dose Emetine against Human Cytomegalovirus. PLoS Pathog. 2016;12(6):e1005717.

65. Chaves Valadao AL, Abreu CM, Dias JZ, Arantes P, Verli H, Tanuri A, et al. Natural Plant Alkaloid (Emetine) Inhibits HIV-1 Replication by Interfering with Reverse Transcriptase Activity. Molecules. 2015;20(6):11474–89.

66. Choy KT, Wong AY, Kaewpreedee P, Sia SF, Chen D, Hui KPY, et al. Remdesivir, lopinavir, emetine, and homoharringtonine inhibit SARS-CoV-2 replication in vitro. Antiviral Res. 2020;178:104786.

67. Khandelwal N, Chander Y, Rawat KD, Riyesh T, Nishanth C, Sharma S, et al. Emetine inhibits replication of RNA and DNA viruses without generating drug-resistant virus variants. Antiviral Res. 2017;144:196–204.

68. Kim JE, Song YJ. Anti-varicella-zoster virus activity of cephalotaxine esters in vitro. J Microbiol. 2019;57(1):74–9.

69. Dong HJ, Wang ZH, Meng W, Li CC, Hu YX, Zhou L, et al. The Natural Compound Homoharringtonine Presents Broad Antiviral Activity In Vitro and In Vivo. Viruses. 2018;10(11).

70. Cao J, Forrest JC, Zhang X. A screen of the NIH Clinical Collection small molecule library identifies potential anti-coronavirus drugs. Antiviral Res. 2015;114:1–10.

71. Romero MR, Serrano MA, Efferth T, Alvarez M, Marin JJ. Effect of cantharidin, cephalotaxine and homoharringtonine on “in vitro” models of hepatitis B virus (HBV) and bovine viral diarrhoea virus (BVDV) replication. Planta Med. 2007;73(6):552–8.

72. Fu Y, Gaelings L, Soderholm S, Belanov S, Nandania J, Nyman TA, et al. JNJ872 inhibits influenza A virus replication without altering cellular antiviral responses. Antiviral Res. 2016;133:23–31.

73. Andreani J, Le Bideau M, Duflot I, Jardot P, Rolland C, Boxberger M, et al. In vitro testing of combined hydroxychloroquine and azithromycin on SARS-CoV-2 shows synergistic effect. Microb Pathog. 2020;145:104228.

74. Sareen, D., Gowing, G., Sahabian, A., Staggenborg, K., Paradis, R., Avalos, P., Latter, J., Ornelas, L., Garcia, L. & Svendsen, C. N. (2014) Human induced pluripotent stem cells are a novel source of neural progenitor cells (iNPCs) that migrate and integrate in the rodent spinal cord, J Comp Neurol. 522, 2707–28.

75. Oberacker, P., Stepper, P., Bond, D. M., Hohn, S., Focken, J., Meyer, V., Schelle, L., Sugrue, V. J., Jeunen, G. J., Moser, T., Hore, S. R., von Meyenn, F., Hipp, K., Hore, T. A. & Jurkowski, T. P. (2019) Bio-On-Magnetic-Beads (BOMB): Open platform for high-throughput nucleic acid extraction and manipulation, PLoS Biol. 17, e3000107.

76. Corman, V. M., Landt, O., Kaiser, M., Molenkamp, R., Meijer, A., Chu, D. K., Bleicker, T., Brunink, S., Schneider, J., Schmidt, M. L., Mulders, D. G., Haagmans, B. L., van der Veer, B., van den Brink, S., Wijsman, L., Goderski, G., Romette, J. L., Ellis, J., Zambon, M., Peiris, M., Goossens, H., Reusken, C., Koopmans, M. P. & Drosten, C. (2020) Detection of 2019 novel coronavirus (2019-nCoV) by real-time RT-PCR, Euro Surveill. 25.

77. Bosl, K., Ianevski, A., Than, T. T., Andersen, P. I., Kuivanen, S., Teppor, M., Zusinaite, E., Dumpis, U., Vitkauskiene, A., Cox, R. J., Kallio-Kokko, H., Bergqvist, A., Tenson, T., Merits, A., Oksenych, V., Bjoras, M., Anthonsen, M., Shum, D., Kaarbo, M., Vapalahti, O., Windisch, M. P., Superti-Furga, G., Snijder, B., Kainov, D. & Kandasamy, R. K. (2019) Common Nodes of Virus–Host Interaction Revealed Through an Integrated Network Analysis, Front Immunology. Accepted.

78. Ianevski, A., Zusinaite, E., Kuivanen, S., Strand, M., Lysvand, H., Teppor, M., Kakkola, L., Paavilainen, H., Laajala, M., Kallio-Kokko, H., Valkonen, M., Kantele, A., Telling, K., Lutsar, I., Letjuka, P., Metelitsa, N., Oksenych, V., Bjoras, M., Nordbo, S. A., Dumpis, U., Vitkauskiene, A., Ohrmalm, C., Bondeson, K., Bergqvist, A., Aittokallio, T., Cox, R. J., Evander, M., Hukkanen, V., Marjomaki, V., Julkunen, I., Vapalahti, O., Tenson, T., Merits, A. & Kainov, D. (2018) Novel activities of safe-in-human broad-spectrum antiviral agents, Antiviral Res. 154, 174–182.

79. Bulanova, D., Ianevski, A., Bugai, A., Akimov, Y., Kuivanen, S., Paavilainen, H., Kakkola, L., Nandania, J., Turunen, L., Ohman, T., Ala-Hongisto, H., Pesonen, H. M., Kuisma, M. S., Honkimaa, A., Walton, E. L., Oksenych, V., Lorey, M. B., Guschin, D., Shim, J., Kim, J., Than, T. T., Chang, S. Y., Hukkanen, V., Kulesskiy, E., Marjomaki, V. S., Julkunen, I., Nyman, T. A., Matikainen, S., Saarela, J. S., Sane, F., Hober, D., Gabriel, G., De Brabander, J. K., Martikainen, M., Windisch, M. P., Min, J. Y., Bruzzone, R., Aittokallio, T., Vaha-Koskela, M., Vapalahti, O., Pulk, A., Velagapudi, V. & Kainov, D. E. (2017) Antiviral Properties of Chemical Inhibitors of Cellular Anti-Apoptotic Bcl-2 Proteins, Viruses. 9.

80. Ianevski, A., He, L., Aittokallio, T. & Tang, J. (2017) SynergyFinder: a web application for analyzing drug combination dose-response matrix data, Bioinformatics. 33, 2413–2415.

81. Desmyter, J., Melnick, J. L. & Rawls, W. E. (1968) Defectiveness of interferon production and of rubella virus interference in a line of African green monkey kidney cells (Vero), J Virol. 2, 955–61.

82. Zhang, Y. N., Zhang, Q. Y., Li, X. D., Xiong, J., Xiao, S. Q., Wang, Z., Zhang, Z. R., Deng, C. L., Yang, X. L., Wei, H. P., Yuan, Z. M., Ye, H. Q. & Zhang, B. (2020) Gemcitabine, lycorine and oxysophoridine inhibit novel coronavirus (SARS-CoV-2) in cell culture, Emerg Microbes Infect, 1–10.

83. Choy, K. T., Wong, A. Y., Kaewpreedee, P., Sia, S. F., Chen, D., Hui, K. P. Y., Chu, D. K. W., Chan, M. C. W., Cheung, P. P., Huang, X., Peiris, M. & Yen, H. L. (2020) Remdesivir, lopinavir, emetine, and homoharringtonine inhibit SARS-CoV-2 replication in vitro, Antiviral Res. 178, 104786.

84. Runfeng, L., Yunlong, H., Jicheng, H., Weiqi, P., Qinhai, M., Yongxia, S., Chufang, L., Jin, Z., Zhenhua, J., Haiming, J., Kui, Z., Shuxiang, H., Jun, D., Xiaobo, L., Xiaotao, H., Lin, W., Nanshan, Z. & Zifeng, Y. (2020) Lianhuaqingwen exerts anti-viral and anti-inflammatory activity against novel coronavirus (SARS-CoV-2), Pharmacol Res. 156, 104761.

85. Fan, H. H., Wang, L. Q., Liu, W. L., An, X. P., Liu, Z. D., He, X. Q., Song, L. H. & Tong, Y. G. (2020) Repurposing of clinically approved drugs for treatment of coronavirus disease 2019 in a 2019-novel coronavirus-related coronavirus model, Chin Med J (Engl). 133, 1051–1056.

86. Cairns, M., Cisse, B., Sokhna, C., Cames, C., Simondon, K., Ba, E. H., Trape, J. F., Gaye, O., Greenwood, B. M. & Milligan, P. J. (2010) Amodiaquine dosage and tolerability for intermittent preventive treatment to prevent malaria in children, Antimicrob Agents Chemother. 54, 1265–74.

87. Schimmer, A. D., Raza, A., Carter, T. H., Claxton, D., Erba, H., DeAngelo, D. J., Tallman, M. S., Goard, C. & Borthakur, G. (2014) A multicenter phase I/II study of obatoclax mesylate administered as a 3- or 24-hour infusion in older patients with previously untreated acute myeloid leukemia, PLoS One. 9, e108694.

88. Mastrangelo, M. J., Grage, T. B., Bellet, R. E. & Weiss, A. J. (1973) A phase I study of emetine hydrochloride (NSC 33669) in solid tumors, Cancer. 31, 1170–5.

89. Bojkova, D., Klann, K., Koch, B., Widera, M., Krause, D., Ciesek, S., Cinatl, J. & Munch, C. (2020) Proteomics of SARS-CoV-2-infected host cells reveals therapy targets, Nature.

