## supplementary information for "Potential antiviral options against SARS-CoV-2 infection"

**Table S1**. Compounds, their suppliers, catalogue numbers and AUCs.

| Drug | CAS | Supplier | pur, % | Cat N | MW | Formula | AUC |
|---|---|---|---|---|---|---|---|
| Abamectin | 71751-41-2 | Dr. Ehrenstorfer GmbH | 95.3 | DRE-CA10001000 | 873 | C95H142O28 | 151,97 |
| Amiloride (EIPA) | 2016-88-8 | Cayman Chemical | >98 | 14409 | 300 | C6H8ClN7O HCl | 182,56 |
| Amiodarone | 1216715-80-8 | Cayman Chemical | >99 | 10010668 | 645 | C25H25D4I2NO3 HCl | 173,88 |
| Amodiaquine | 86-42-0 | Cayman Chemical | >95 | 15954 | 356 | C20H22ClN3O | 404,88 |
| Apilimod | 54550-10-0 | Axon MedChem | 99.7 | 1369 | 419 | C23H26N6O2 | 585,46 |
| Arbidol | 131707-23-8 | Cayman Chemical | >98.5 | 16933 | 477 | C22H25BrN2O3S HCl | 319,27 |
| Artesunate | 88495-63-0 | Acros | n/a | 460340050 | 384 | C19H28O8 | 222,25 |
| BDA-366 | 1909226-00-1 | Selleckchem | n/a | S7849 | 423,5 | C24H29N3O4 | 174,3 |
| Bepridil | 68099-86-5 | Cayman Chemical | >98 | 19645 | 403 | C24H34N2O HCl | 285,95 |
| Berberine | 633-65-8 | Cayman Chemical | >95 | 10006427 | 336 | C20H18ClNO4 | 236,67 |
| Brequinar | 96187-53-0 | Cayman Chemical | >98 | 24445 | 375 | C23H15F2NO2 | 260,54 |
| Bromocriptine | 22260-51-1 | Cayman Chemical | >98 | 14598 | 656 | C32H40BrN5O5 CH3SO3H | 164,57 |
| Camostat | 59721-29-8 | Cayman Chemical | >98 | 16018 | 398 | C20H22N4O5 CH3SO3H | 88,41 |
| Camptothecin | 7689-03-4' | Cayman Chemical | >98 | 11694 | 348 | C20H16N2O4 | 57,75 |
| DFMO (Difluoromethylomithine) | 70052-12-9 | ChemCruz | >98 | sc-204723 | 182 | C6H12F2N2O2 | 142,66 |
| Diphylline (7-(2,3-Dihydroxypropyl)theophylline) | 479-18-5 | Acros | 99 | 115051000 | 524 | C10H14N4O4 | 195,44 |
| Emetine | 7083-71-8 | Cayman Chemical | >98 | 21048 | 481 | C29H40N2O4 2HCl | 484,19 |
| Emodin | 518-82-1 | Cayman Chemical | >98 | 13109 | 270 | C15H10O5 | 170,24 |
| Fenretinide (4-HPR) | 65646-68-6 | Cayman Chemical | >98 | 17688 | 392 | C26H33NO2 | 137,9 |
| Formoterol | 183814-30-4 | Sigma-Aldrich | n/a | F0372000 | 420 | C19H24N2O4 0.5C4H4O4 H2O | 203,77 |
| Ganciclovir | 82410-32-0 | Sigma-Aldrich | >99 | G2536-100MG | 255,23 | C9H13N5O4 | 243,88 |
| Glycyrrhizic acid (Glycyrrhizin) | 1405-86-3 | MCE | >98 | HY-N0184 | 823 | C42H62O16 | 235,76 |
| Homoharringtonine | 26833-87-4 | Cayman Chemical | >98 | 14631 | 546 | C29H39NO9 | 374,71 |
| Kasugamycin | 19408-46-9 | Fluka | >96.5 | R1114-100MG | 379 | C14H25N3O9 | 88,55 |
| Lanatoside C | 17575-22-3 | MCE | 99 | HY-B1030 | 985 | C49H76O20 | 54,25 |
| Letermovir | 917389-32-3 | Cayman Chemical | >98 | 17556 | 573 | C29H28F4N4O4 | 134,12 |
| Lobucavir | 127759-89-1 | Santa Cruz | 98 | sc-211744 | 265 | C11H15N5O3 | 153,65 |
| Luteolin | 491-70-3 | Santa Cruz | >98 | sc-203119 | 286 | C15H10O6 | 119,07 |
| Manidipine | 89226-50-6 | Cayman Chemical | >98 | 23614 | 611 | C35H38N4O6 | 133 |
| Maribavir | 176161-24-3 | MCE | >98 | HY-16305 | 376 | C15H19Cl2N3O4 | 145,81 |
| Mitoxantrone | 70476-82-3 | Cayman Chemical | 95 | 14842 | 517 | C22H28N4O6 HCl | 125,09 |
| Nafamostat | 82956-11-4 | Cayman Chemical | >98 | 14837 | 347 | C19H17N5O2 2CH3SO3H | 166,53 |

| | | | | | | | |
|---|---|---|---|---|---|---|---|
| Nelfinavir | 159989-65-8 | Cayman Chemical | >98 | 15144 | 664 | C32H45N3O4S CH3SO3H | 286,37 |
| Novobiocin | 303-81-1 | Carbosynth | >94 | FN59697 | 612 | C31H36N2O11 | 100,73 |
| Niclosamide | 50-65-7 | Dr. Ehrenstorfer GmbH | 97 | C15510000 | 327 | C13H8Cl2N2O4 | 149,52 |
| Obatoclax (Mesylate) | 803712-79-0 | MedChemExpress | 99,7 | HY-10969/CS-0133 | 413,49 | $C_{21}H_{23}N_3O_4S$ | 385,91 |
| Posaconazole | 171228-49-2 | Cayman Chemical | >95 | 14737 | 701 | C37H42F2N8O4 | 318,22 |
| PSI-7977 (Sofosbuvir) | 1190307-88-0 | MCE | 100 | HY-15005 | 530 | C22H29FN3O9P | 131,11 |
| Quinine | 130-95-0 | Alfa Aesar | 99 | A10459.09 | 324 | C20H24N2O2 | 140,56 |
| Raloxifene | 82640-04-8 | Cayman Chemical | >98 | 10011620 | 474 | C28H27NO4S HCl | 199,57 |
| Regorafenib | 755037-03-7 | Cayman Chemical | >98 | 18498 | 483 | C21H15ClF4N4O3 | 24,12 |
| Roscovitine | 186692-46-6 | Cayman Chemical | >98 | 10009569 | 355 | C19H26N6O | 137,97 |
| Saikosaponin A | 20736-09-8 | Sigma Aldrich | n/a | Y0001932 | 780,98 | C42H68O13 | 77 |
| Sorafenib (BAY 43-9006) | 284461-73-0 | Cayman Chemical | >98 | 10009644 | 465 | C21H16ClF3N4O3 | 33,12 |
| Suramin | 129-46-4 | Acros | 98 | 328540500 | 1297 | C51H40N6O23S6 | 94,57 |
| Tamoxifen | 10540-29-1 | Sigma Aldrich | >99 | 85256-50MG | 372 | C26H29NO | 54,67 |
| Zanamivir | 139110-80-8 | Cayman Chemical | ≥98 | 15123 | 332,3 | C12H20N4O7 | 113,54 |
| Genistin | 529-59-9 | Cayman Chemical | ≥98 | 14174 | 432,4 | C21H20O10 | 54,1 |
| Pleconaril | 153168-05-9 | Cayman Chemical | ≥98 | 28461 | 381 | C18H18F3N3O3 | 141,75 |
| Salinomycin | 53003-10-4 | MedChem Express | ≥98 | HY-15597 | 751 | C42H70O11 | 334,04 |
| Monensin | 22373-78-0 | Cayman Chemical | ≥98 | 16488 | 671 | C36H61O11 • Na | 254,31 |
| Enoxacin | 74011-58-8 | Alfa Aesar | n/a | J61912 | 320 | C15H17FN4O3 | 118,23 |
| Hexachlorophene | 70-30-4 | Cayman Chemical | ≥98 | CAYM23948 | 407 | C13H6Cl6O2 | 122,15 |
| SNS-032 | 345627-80-7 | Selleckchem | ≥98 | S1145 | 380,5 | C17H24N4O2S2 | 96,25 |
| Trametinib | 871700-17-3 | MedChemExpress | 9944 | HY-10999/CS-0060 | 615,39 | $C_{26}H_{23}FIN_5O_4$ | 117,53 |
| Ciclesonide | 126544-47-6 | Sigma-Aldrich | 98 | SML1955-5MG | 540,69 | C32H44O7 | 157,15 |
| Pimodivir | 1629869-44-8 | MedChemExpress | 99 | HY-12353A/CS | 399,39 | $C_{20}H_{19}F_2N_5O_2$ | 126,91 |
| Ouabain octahydrate | 11018-89-6 | Sigma-Aldrich | 95 | O3125-250MG | 728,77 | C29H44O12・8H2O | 41,93 |
| Tilorone (dihydrochloride) | 27591-69-1 | MedChemExpress | 99 | HY-B1080/CS-4636 | 483,47 | $C_{25}H_{36}Cl_2N_2O_3$ | 83,72 |
| Staurosporine | 62996-74-1 | Sigma-Aldrich | 98 | S4400-1MG | 466,53 | C22H18O12 | 15,75 |
| Chicoric acid | 6537-80-0 | Sigma-Aldrich | 95 | C7243-10MG | 474,37 | C22H18O12 | 93,1 |
| Ascorbic acid | 50-81-7 | Sigma-Aldrich | 98 | A4544-25g | 176,12 | C6H8O6 | 108,5 |
| Acetylsalicylic acid | 50-78-2 | Acros Organics | 99 | 158180500 | 180 | C9H8O4 | 131,04 |
| Aciclovir (acycloguanosine) | 59277-89-3 | Acros Organics | 98 | 445240100 | 225 | C8H11N5O3 | 124,95 |
| Azacytidine | 320-67-2 | Acros Organics | 99 | 226620500 | 244 | C8H12N4O5 | 228,06 |
| Azithromycin | 83905-01-5 | Santa Cruz Biotechnology | >95 | SC-254949 | 749 | C38H72N2O12 | 119,42 |
| BCX4430 (galidesivir) | 249503-25-1 | MedChem Express | >99 | HY-18649 | 265 | C11H15N5O3 | 94,08 |

| | | | | | | | |
|---|---|---|---|---|---|---|---|
| Bortezomib (PS-341) | 179324-69-7 | Selleckchem | 99 | S1013 | 384 | C19H25BN4O4 | 68,67 |
| Brincidofovir (CMX001) | 444805-28-1 | MedChem Express | >98 | HY-14532 | 562 | C27H52N3O7P | 126,49 |
| Caffeine | 58-08-2 | Acros Organics | >98 | 108160100 | 194 | C8H10N4O2 | 147,28 |
| Chloroquine phosphate | 50-63-5 | Sigma-Aldrich | >99 | PHR1258 | 516 | C18H26ClN3 2H3PO4 | 165,9 |
| Cidofovir | 113852-37-2 | Cayman Chemical | 95 | CAYM13113 | 279 | C8H14N3O6P | 144,69 |
| Clofarabine | 123318-82-1 | Sigma-Aldrich | 98 | C7495-5MG | 303,68 | C10H11ClFN5O3 | 112,42 |
| Cyclosporine A | 59865-13-3 | Acros Organics | 98 | 457970010 | 1202 | C62H111N11O12 | 146,37 |
| Dasatinib | 302962-49-8 | Sigma-Aldrich | >99 | CDS023389 | 488 | C22H26ClN7O2S | 135,45 |
| Dibucaine | 61-12-1 | Sigma-Aldrich | 99 | D0638 | 380 | C20H29N3O2 HCl | 214,34 |
| Doxycycline hyclate | 24390-14-5 | Sigma-Aldrich | >98 | D9891 | 513 | C22H24N2O8 HCl 0.5H2O 0.5C2H6O | 143,99 |
| Erlotinib | 183319-69-9 | Sigma-Aldrich | >99 | CDS022564 | 430 | C22H24ClN3O4 | 103,46 |
| Ezetimibe | 163222-33-1 | Cayman Chemical | >98 | CAYM16331 | 409 | C24H21F2NO3 | 138,39 |
| Esomeprazole | 668985-31-7 | Sigma-Aldrich | >98 | E7906 | 713 | C34H36MgN6O6S2 H2O | 177,52 |
| Famciclovir | 104227-87-4 | Sigma-Aldrich | >98 | F7932 | 321 | C14H19N5O4 | 149,52 |
| Favipiravir (T-705) | 36791-04-5 | Selleckchem | 98 | S7975 | 244 | C8H12N4O5 | 136,64 |
| Fluoxetine | 56296-78-7 | Santa Cruz Biotechnology | >98 | SC-201125 | 345 | C17H18F3NO HCl | 104,37 |
| Fluvastatin | 93957-55-2 | Acros Organics | >97 | 458010010 | 433 | C24H25FNNaO4 | 154,14 |
| Foscarnet | 63585-09-1 | Selleckchem | >98 | S3076 | 192 | CNa3O5P | 127,26 |
| Gefitinib | 184475-35-2 | Santa Cruz Biotechnology | >99 | sc-202166 | 447 | C22H24ClFN4O3 | 188,86 |
| Gemcitabine | 122111-03-9 | Sigma-Aldrich | 98 | G6423 | 300 | C9H11F2N3O4 · HCl | 92,89 |
| Hydroxychloroquine | 747-36-4 | Santa Cruz Biotechnology | >97 | sc-215157 | 434 | C18H26ClN3O•H2SO4 | 147,63 |
| Imatinib | 220127-57-1 | Sigma-Aldrich | >98 | SML1027 | 590 | C29H31N7O · CH4O3S | 177,8 |
| Indomethacin | 53-86-1 | Acros Organics | >97 | 458030050 | 358 | C19H16ClNO4 | 104,51 |
| Itraconazole | 84625-61-6 | Acros Organics | 99 | 452870050 | 706 | C35H38Cl2N8O4 | 110,18 |
| Ivermectin | 70288-86-7 | Alfa Aesar | >90 | J62777 | 875 | C48H74O14 | 66,08 |
| Lamivudine | 134678-17-4 | Selleckchem | >99 | S1706 | 229 | C8H11N3O3S | 106,75 |
| Leflunomide | 75706-12-6 | Santa Cruz Biotechnology | >99 | SC-202209 | 270 | C12H9F3N2O2 | 47,95 |
| Lopinavir | 192725-17-0 | Sigma-Aldrich | >98 | SML1222 | 629 | C37H48N4O5 | 157,36 |
| Lovastatin | 75330-75-5 | Sigma-Aldrich | >99 | PHR1285 | 405 | C24H36O5 | 204,26 |
| Memantine | 41100-52-1 | Acros Organics | 99 | 298080010 | 216 | C12H21N HCl | 115,78 |
| Metformin (1,1-Dimethylbiguanide) | 214-230-6 | Alfa Aesar | >97 | J63361 | 166 | C4H11N5 HCl | 116,76 |
| Minocycline | 13614-98-7 | Cayman Chemical | >98 | CAYM14454 | 458 | C23H27N3O7 HCl | 119,14 |
| Mycophenolic acid | 24280-93-1 | Acros Organics | 98 | 459380010 | 320 | C17 H20 O6 | 103,74 |
| Nitazoxanide | 55981-09-4 | Cayman Chemical | >95 | CAYM13692 | 307 | C12H9N3O5S | 120,33 |
| Omeprazole | 73590-58-6 | Sigma-Aldrich | >99 | O104 | 345 | C17H19N3O3S | 139,02 |
| Oritavancin | 192564-14-0 | Sigma-Aldrich | >97 | SML1586 | 1989 | C86H97Cl3N10O26 2H3PO4 | 163,24 |
| Pentosan polysulfate | 9062-57-1 | MedChem Express | >98 | HY-A0203 | 602 | C10H18O21S4 | 123,06 |
| Pirlindole mesylate | 60762-57-4 | Santa Cruz Biotechnology | >99 | SC-203664 | 322 | C15H18N2 CH3SO3H | 120,4 |

| | | | | | | | |
|---|---|---|---|---|---|---|---|
| Rapamycin | 53123-88-9 | Fisher scientific | >98 | BP2963 | 914 | C51H79NO13 | 46,9 |
| Ribavirin | 36791-04-5 | Acros Organics | 98 | 460480010 | 244 | C8H12N4O5 | 134,26 |
| Ritonavir | 155213-67-5 | Cayman chemical co | >98 | CAYM13872 | 721 | C37H48N6O5S2 | 137,69 |
| Simvastatin | 79902-63-9 | Acros Organics | 98 | 458840010 | 419 | C25H38O5 | 163,03 |
| Teicoplanin | 61036-62-2 | Sigma-Aldrich | >99 | Y0001102 | 1879 | Variable | 145,39 |
| Telavancin | 372151-71-8 | Adooq Biosc | >98 | A12698 | 1755 | C80H106Cl2N11O27P | 139,65 |
| Tenofovir disoproxil fumarate | 202138-50-9 | Acros Organics | 98 | 461250010 | 636 | C19H30N5O10P C4H4O4 | 137,2 |
| Topotecan | 119413-54-6 | Santa Cruz Biotechnology | 99 | SC-204919 | 458 | C23H23N3O5 HCl | 124,6 |
| Trifluridine | 70-00-8 | Selleckchem | >99 | S1778 | 296 | C10H11F3N2O5 | 107,17 |
| Valaciclovir | 124832-27-5 | Sigma-Aldrich | >99 | Y0001225 | 361 | C13H20N6O4 HCl | 134,61 |
| Verapamil | 152-11-4 | Acros Organics | 99 | 329330010 | 491 | C27H38N2O4 HCl | 134,26 |
| Bromhexine HCl | 611-75-6 | Sigma-Aldrich | 98 | 17343-25G | 412,59 | C14H20Br2N2 · HCl | 124,95 |
| Bromcriptine mesilate | 22260-51-1 | Sigma-Aldrich | n/a | Y0000677 | 750,7 | C32H40BrN5O5 · CH4SO3 | 59,71 |
| Cefoperazone acid | 62893-19-0 | Santa Cruz Bio | >98 | SC-204677 | 545,7 | C25H27N9O8S2 | 22,05 |
| Moxalactam sodium salt | 64963-12-4 | Sigma-Aldrich | n/a | M8158-1G | 564,4 | C25H27N9O8S2 | 18,13 |
| Anisomycin | 22862-76-6 | Sigma-Aldrich | >98 | A9789-5mg | 265,3 | C14H19NO4 | 169,33 |
| Benzatropine | 132-17-2 | Sigma-Aldrich | >98 | SML0847-500mg | 403,53 | C21H25NO · CH3SO3H | 208,11 |
| Diltiazem | 33286-22-5 | Sigma-Aldrich | n/a | D1980000 | 414,519 | C22H26N2O4S | 99,61 |
| Teriflunomide | 15345-81-8 | Sigma-Aldrich | >98 | SML0936-10MG | 270,2 | C12H9F3N2O2 | 72,87 |
| Clomiprade hydrochloride | 17321-77-6 | Sigma-Aldrich | >98 | C7291-1G | 351,31 | C19H23ClN2 · HCl | 74,69 |
| Chlorpromazine hydrochloride | 69-09-0 | Sigma-Aldrich | >98 | C8138-5G | 356,33 | C17H19ClN2S · HCl | 86,52 |
| Quinacrine dihydrochloride | 69-05-6 | Sigma-Aldrich | >90 | Q3251-25G | 472,88 | C23H30ClN3O · 2HCl | 105,63 |
| Toremifene | 89778-27-8 | Sigma-Aldrich | >98 | T7204-5MG | 598,1 | C32H36ClNO8 | 27,02 |
| Ranitidine | 66357-35-5 | Sigma-Aldrich | n.a. | R101-1G | 350,9 | C13H22N4O3S · HCl | 94,08 |
| Stanozolol | 10418-03-8 | Sigma-Aldrich | n.a. | S1250000 | 328,5 | C21H32N2O | 161,7 |
| Bumetanide | 28395-03-1 | Sigma-Aldrich | >98 | B3023-250MG | 364,4 | C17H20N2O5S | 87,85 |
| Dalbavancin | 171500-79-1 | Sigma-Aldrich | >98 | SML2378-5MG | 1816 | C88H101Cl3N10O28 | 136,29 |
| Remdesivir | 1809249-37-3 | Cayman | >98 | 30354 | 602,6 | C27H35N6O8P | 290,29 |
| Trametinib | 871700-17-3 | Cayman | >98 | 16292 | 615,4 | C26H23FIN5O4 | 100,59 |
| Acetylcysteine | 616-91-1 | Cayman | >98 | 20261 | 163,2 | C5H9NO3S | 43,47 |
| Verdinexor | 1392136-43-4 | Cayman | >98 | 26171 | 442,3 | C18H12F6N6O | 83,93 |
| Mefloquine | 51773-92-3 | Sigma-Aldrich | n.a. | M0253000 | 414,8 | C17H17ClF6N2O | 189.28 |

**Table S2**. Results of neutralization and ELISA assays.

| Sample N | s/co, IgM | s/co, IgG | SSS | Virus |
|---|---|---|---|---|
| 10 | 14,35 | 2,11 | 17,8 | SARS-CoV-2 |
| 11 | 2,25 | 2,92 | 37,6 | SARS-CoV-2 |
| 12 | 3,59 | 2,95 | 5 | SARS-CoV-2 |
| 13 | 1,44 | 3,02 | 16,1 | SARS-CoV-2 |

| | | | | |
|---|---|---|---|---|
| 14 | 0,48 | 0,26 | 6,1 | SARS-CoV-2 |
| 15 | 0,53 | 3,77 | 2,5 | SARS-CoV-2 |
| 16 | 0,43 | 0,39 | 0,9 | SARS-CoV-2 |
| 17 | 0,43 | 2,56 | 17,1 | SARS-CoV-2 |
| 18 | 0,53 | 1,10 | 5,7 | SARS-CoV-2 |
| 19 | 0,48 | 0,55 | 0,3 | SARS-CoV-2 |
| 26 | 0,43 | 0,94 | 0 | SARS-CoV-2 |
| 27 | 0,48 | 1,33 | 1,8 | SARS-CoV-2 |
| 28 | 0,91 | 3,02 | 7,4 | SARS-CoV-2 |
| 29 | 0,41 | 1,33 | 0 | SARS-CoV-2 |
| 30 | 0,41 | 2,05 | 0,8 | SARS-CoV-2 |
| 31 | 0,48 | 1,85 | 2,6 | SARS-CoV-2 |
| 32 | 0,62 | 2,31 | 2,9 | SARS-CoV-2 |
| 81 | 0,53 | 0,52 | 0 | HCoV-HKU1 |
| 82 | 0,48 | 0,58 | 0,5 | HCoV-NL63 |
| 83 | 0,53 | 0,34 | 0 | HCoV-NL63 |
| 86 | 0,48 | 0,48 | 0,5 | HCoV-OC43 |
| 88 | 0,48 | 0,75 | 0,4 | HCoV-OC43 |
| 91 | 0,48 | 0,61 | 2,6 | HCoV-229E |
| 92 | 1,20 | 0,51 | 1,4 | HCoV-229E |
| 72 | 0,35 | 0,27 | 0 | Healthy blood donor |
| 73 | 0,35 | 0,45 | 0 | Healthy blood donor |
| 74 | 0,39 | 0,45 | 0 | Healthy blood donor |
| 75 | 0,39 | 0,55 | 0 | Healthy blood donor |
| 76 | 0,39 | 0,30 | 0 | Healthy blood donor |
| 77 | 0,43 | 0,18 | 0 | Healthy blood donor |
| 78 | 0,39 | 0,36 | 0 | Healthy blood donor |
| 79 | 0,35 | 0,85 | 0 | Healthy blood donor |

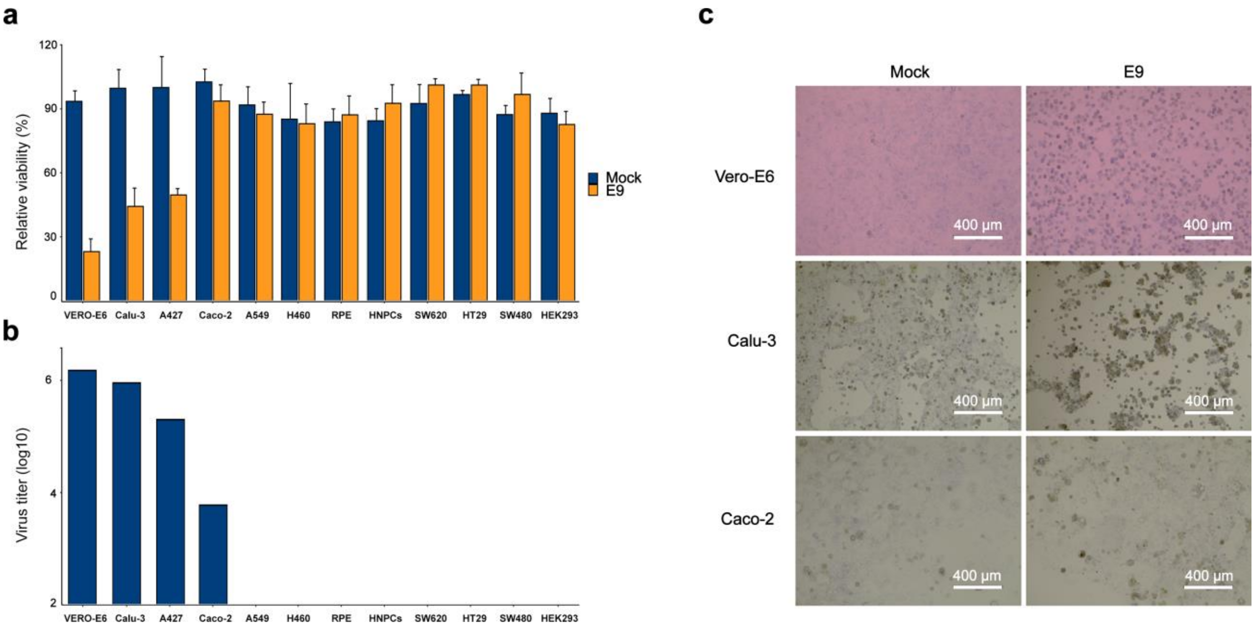


**Figure S1.** Propagation of HCoV-19/Norway/Trondheim-E9/2020 in cell cultures. (a) Cell lines were infected with the virus and cell viability were measured. (b) Viruses amplified in the cells were quantified by plaque assay. (c) Viability of mock- and virus-infected cells were visualized by microscopy (bright field).

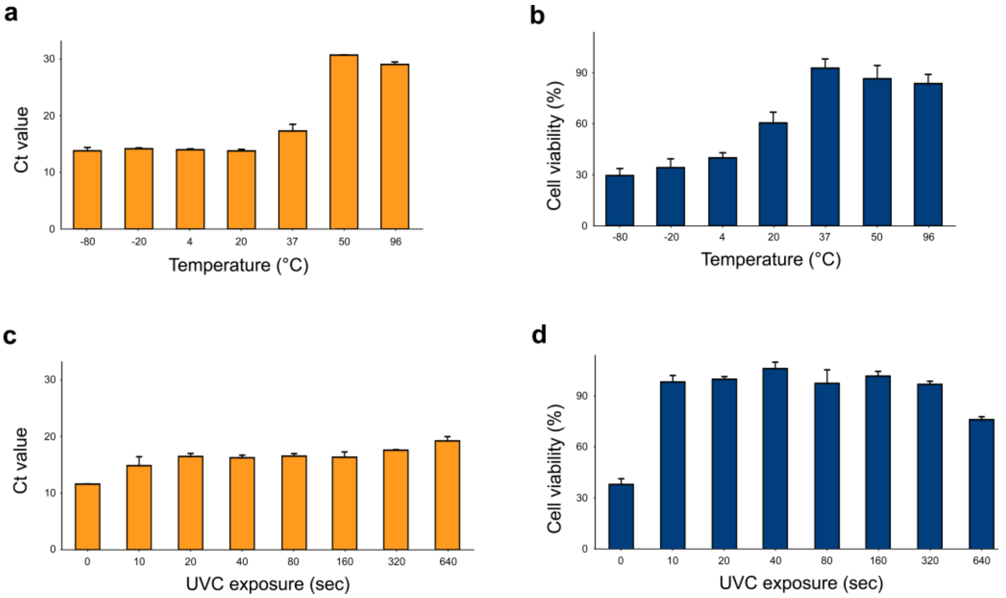


**Figure S2.** Effect of temperature and UV radiation on infectivity of HCoV-19/Norway/Trondheim-E9/2020 strain. (a) The virus was incubated at -80, -20, 4, 20, 37, and 50 °C for 48 h or at 96 °C for 10 min. The thermostability of viral RNA was analysed by RT-qPCR. (b) Vero-E6 cells were subsequently infected with the virus. After 72 h cell viability was measured. Mean ± SD, n = 3. (c) The virus was exposed to UVC for different times. The stability of viral RNA was analysed by RT-qPCR. Mean ± SD, n = 3. (d) Vero-E6 cells were subsequently infected with the virus. After 72 h cell viability was measured. Mean ± SD, n = 3.

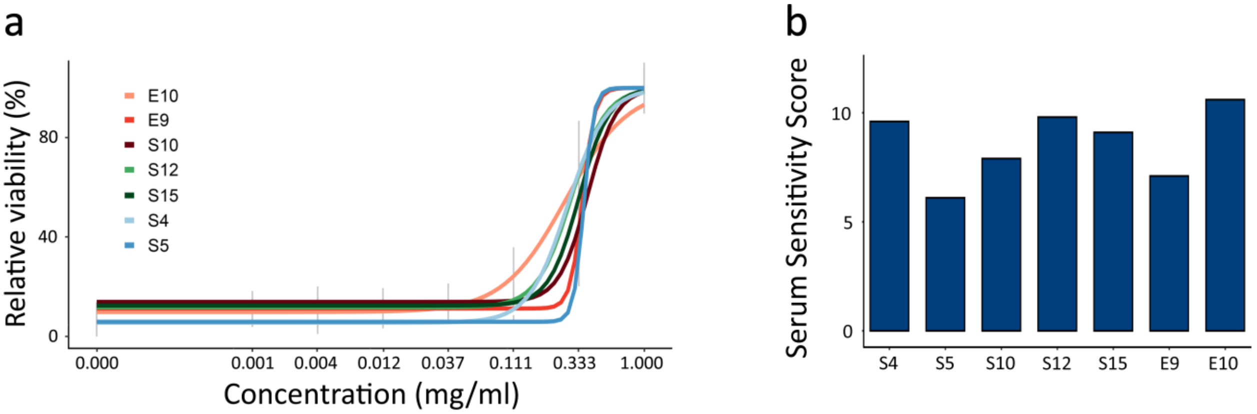


**Figure S3**. The effect of serum from patient recovered from COVID-19 on viability of Vero-E6 cells infected with 7 SARS-CoV-2 strains.

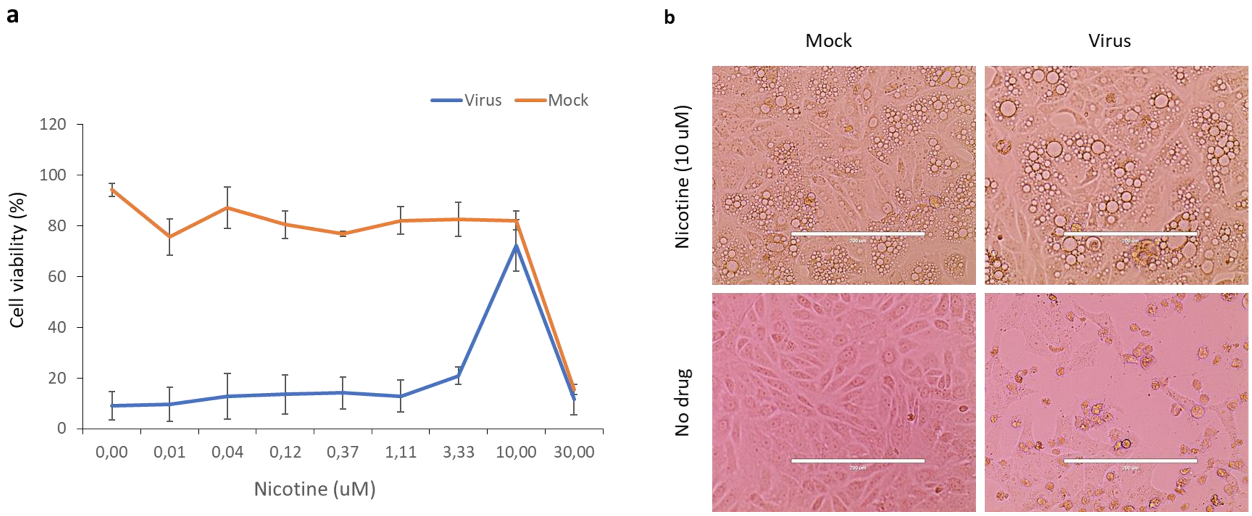


**Figure S4**. The effect of nicotine on viability and morphology of mock- and SARS-CoV-2-infected Vero-E6 cells. (**a**) Cells were treated with increasing concentrations of a compound and infected with HCoV-19/Norway/Trondheim-E9/2020 strain (moi, 0.1) or mock. After 72 h the viability of the cells was determined using the CTG assay. Mean ± SD; n = 3. (**b**) Cells were treated as for (a). 72 h after infection cells were imaged using a bright-field microscopy. Scale bar, 200 µm.

**a**

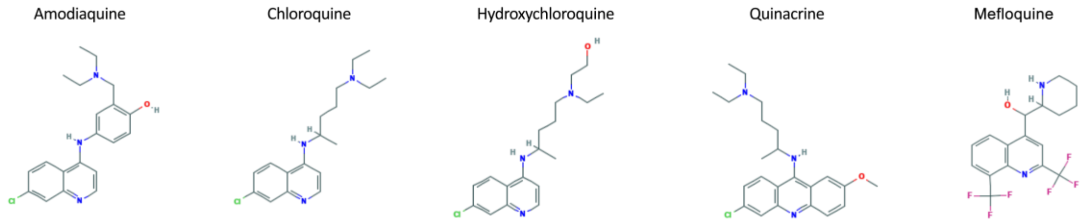


**b**

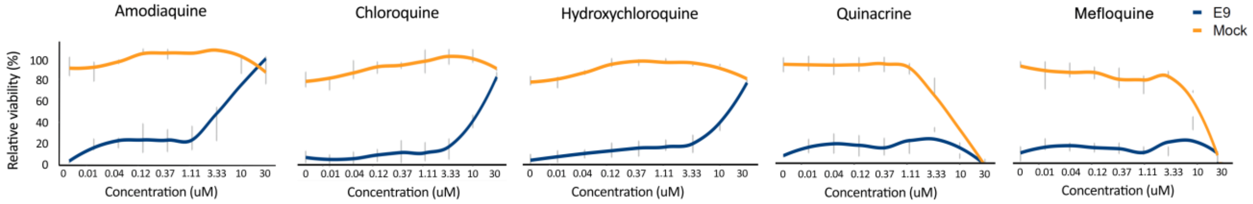


**c**

| Drug | CC50, uM | EC50_1, uM | EC50_2, uM | SI_1 | SI_2 |
|---|---|---|---|---|---|
| Amodiaquine | >30 | 4,2±0,2 | 5,0±0,7 | >7,1 | >6,0 |
| Chloroquine | >30 | 15,5±1,0 | 18,0±1,6 | >1,9 | >1,7 |
| Hydroxychloroquine | >30 | 13,7±0,8 | 16,2±1,2 | >2,2 | >1,9 |
| Quinacrine | 13,9±0,3 | n.a. | 8,4±2,3 | n.a. | 1,7 |
| Mefloquine | 16.1±0,1 | n.a. | n.a. | n.a. | n.a. |

**Figure S5**. Comparison of anti-SARS-CoV-2 activities of amodiaquine and its analogues. (**a**) Structure of amodiaquine and its analogues. (**b**) Anti-SARS-CoV-2 activity of amodiaquine and its analogues in Vero-E6 cells. Cells were treated with increasing concentrations of a compound and infected with HCoV-19/Norway/Trondheim-E9/2020 strain (moi, 0.1) or mock. After 72 h the viability of the cells was determined using the CTG assay. Mean ± SD; n = 3. (**c**) Table showing half-maximal cytotoxic concentration ($CC_{50}$), the half-maximal effective concentration ($EC_{50}$), and selectivity indexes ($SI=CC_{50}/EC_{50}$) for amodiaquine and its analogues calculated from CTG and plaque assays. Mean ± SD; n = 3.
